# Serial Capture Affinity Purification and Integrated Structural Modeling of the H3K4me3 Binding and DNA Damage Related WDR76:SPIN1 Complex

**DOI:** 10.1101/2023.01.31.526478

**Authors:** Xingyu Liu, Ying Zhang, Zhihui Wen, Yan Hao, Charles A.S. Banks, Jeffrey J. Lange, Joseph Cesare, Saikat Bhattacharya, Brian D. Slaughter, Jay R. Unruh, Laurence Florens, Jerry L. Workman, Michael P. Washburn

## Abstract

WDR76 is a multifunctional protein involved in many cellular functions. With a diverse and complicated protein interaction network, dissecting the structure and function of specific WDR76 complexes is needed. We previously demonstrated the ability of the Serial Capture Affinity Purification (SCAP) method to isolate specific complexes by introducing two proteins of interest as baits at the same time. Here, we applied SCAP to dissect a subpopulation of WDR76 in complex with SPIN1, a histone marker reader that specifically recognizes trimethylated histone H3 lysine4 (H3K4me3). In contrast to the SCAP analysis of the SPIN1:SPINDOC complex, H3K4me3 was copurified with the WDR76:SPIN1 complex. In combination with crosslinking mass spectrometry, we built an integrated structural model of the complex which revealed that SPIN1 recognized the H3K4me3 epigenetic mark while interacting with WDR76. Lastly, interaction network analysis of copurifying proteins revealed the potential role of the WDR76:SPIN1 complex in the DNA damage response.

**Teaser:** In contrast to the SPINDOC/SPIN1 complex, analyses reveal that the WDR76/SPIN1 complex interacts with core histones and is involved in DNA damage.

## Introduction

The human genome project has largely increased the annotation of the human proteome (*2*). Subsequently, the functional investigation of newly identified proteins has become a challenging task. In cells, most proteins do not function in isolation. Thus, one key approach to unraveling the biological function of proteins is to identify protein-protein interactions (PPIs). Affinity purification coupled with mass spectrometry (AP-MS) has been used widely to identify protein complex interactions (*3*), even in studies of large-scale human PPI networks (*4–7*). Despite of many advantages of AP-MS, there are several common challenges to interpret the data, including distinguishing *bona fide* interactions from contaminants and resolving the subsets of interactors when the bait protein is multifunctional or associated with multiple complexes.

We came across those challenges when investigating the function of a largely unknown WD40 repeat containing protein, wdr76 (CMR1 or Ydl156w), identified in *S. cerevisiae* (*8*). Previous AP-MS analyses of human WDR76 from our lab have identified a number of intriguing protein interactions that alluded to distinct functions (*9, 10*) (**Fig. 1A**). Evidence from other studies also indicate the roles of WDR76 in diverse biological activities. Studies in different model organisms have discovered WDR76 was involved in DDR (*8, 9, 11-14*). In addition to a potential function in DDR, studies have uncovered that WDR76 might also play a role in transcriptional regulation (*8, 15–17*). Moreover, WDR76 has also been involved in ubiquitination activities (*18–22*). Although some of the activities mentioned above might have overlaps, it is unlikely that all legitimate interactions of WDR76 are functionally related in cells. To try and parse out the precise role of WDR76, we sought to target a subpopulation of WDR76-containing complexes.

**Fig. 1.**
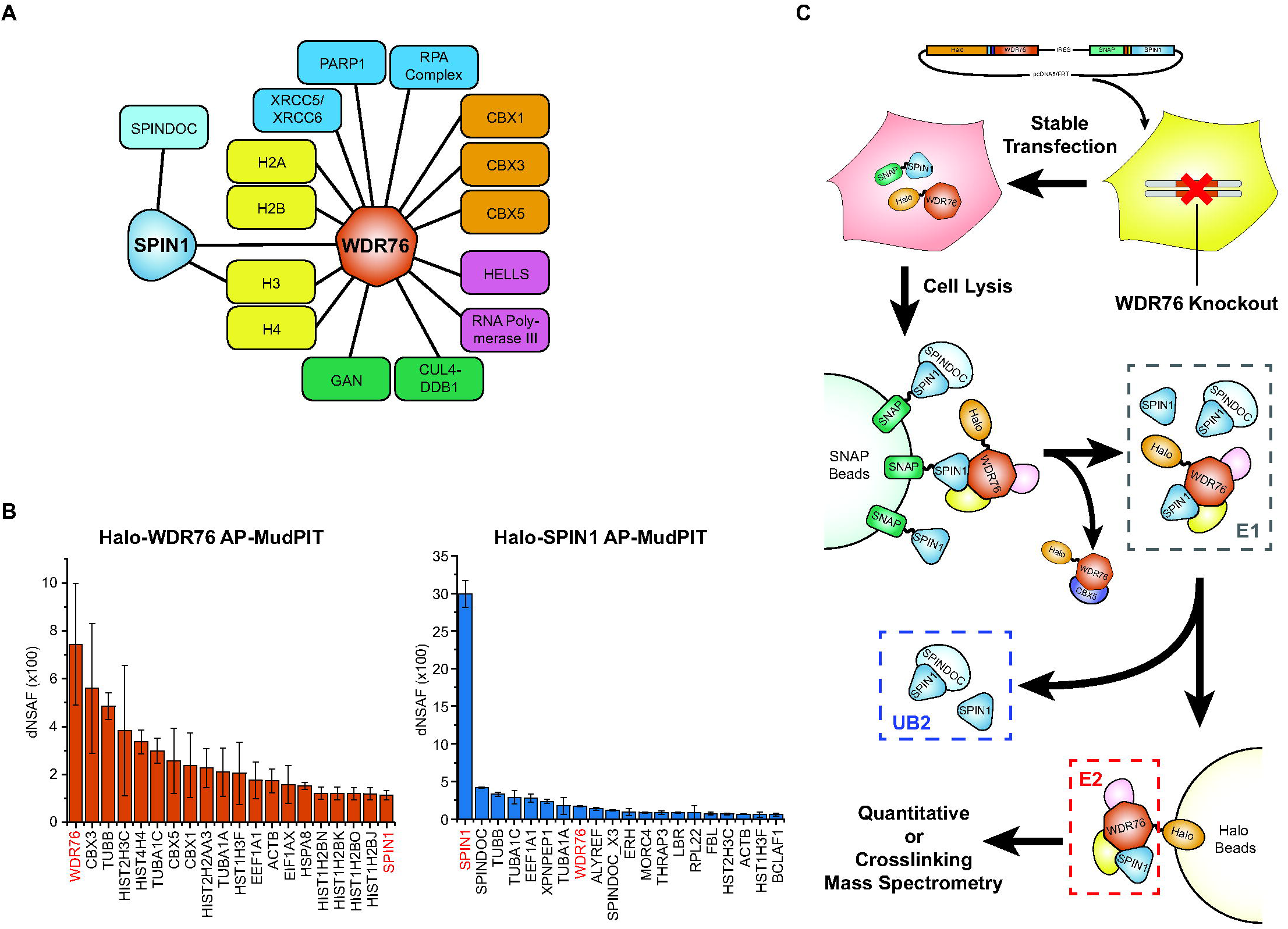
Modified Serial Capture Affinity Purification (SCAP) method to study WDR76 and SPIN1 containing protein complex. (**A**) Selected potential interacting proteins of WDR76 and SPIN1 identified by affinity purification coupled with mass spectrometry (AP-MS) (*8–10*). (**B**) MudPIT analysis of WDR76 co-purified and SPIN1 co-purified proteins. Top 20 proteins ranked by dNSAF values in Halo-WDR76 (left) or Halo-SPIN1 (right) purification are showed. Each dNSAF value plotted is the average from three biological replicates; error bars represent standard deviations. (**C**) Schema of using SCAP method to investigate co-interacting proteins of WDR76 and SPIN1. The modified SCAP pipeline started with generating the stable cell line expressing tagged bait proteins in absent of endogenous version of the second bait protein (WDR76 in this example), and then this cell line was used to provide material for SNAP purification followed by Halo purification. Samples might be analyzed by quantitative mass spectrometry (SCAP-MS) or be subjected to crosslinking steps (SCAP-XL).

An important component of AP-MS identified WDR76 co-purified proteins is the core histone subunits (*8, 9*) (**Fig. 1B**), indicating its involvement in chromatin related activities. Intriguingly, another WDR76 co-purified protein, Spindlin-1 (SPIN1 or ORC) is also a histone binding protein. SPIN1 contains Tudor domains and has been proved to specifically bind H3K4me3 containing histone H3 peptide with high affinity (*23–26*). The interaction between WDR76 and SPIN1 is potentially significant to human health as SPIN1 is reported to promote cancer cell proliferation in multiple types of cancers (*27–29*). However, characterizing the complex of WDR76/SPIN1 and other co-associated proteins could be challenging using conventional AP-MS approaches. Key interactions of WDR76 and SPIN1 reported by others and our group were summarized in **Figure 1A**. Among the most abundant WDR76 co-purified proteins, the chromobox proteins (CBX1, CBX3, CBX5) are mammalian orthologues of the *Drosophila* heterochromatin protein 1(HP1) (**Fig. 1B**). They are not likely to be in the same complex with SPIN1 as they recognize opposite epigenetic markers (*30–32*). Likewise, SPINDOC (C11orf84), the most abundant interacting protein of SPIN1 (**Fig. 1B**), probably forms a different complex than SPIN1 forms with WDR76, as SPINDOC was not co-purified with WDR76 (*9, 10*) and it prevents SPIN1 from binding H3K4me3 (*1, 33, 34*). Moreover, despite that histone H3 was co-purified with both WDR76 and SPIN1, there is no sufficient evidence showing whether they interact with H3 as a complex or these are independent interactions. To this point, purification methods that effectively enrich a specific subset of interactions are keenly needed for characterize protein complexes formed by proteins engaged in multiple protein complexes, like WDR76 and SPIN1.

In our previous publication, we have described Serial Capture Affinity Purification (SCAP), in which a combination of two separately tagged bait proteins, Halo-tagged SPIN1 and SNAP-tagged SPINDOC, were co-expressed and purified in sequence to reduce the complexity of purified samples (*1*). In the same study, we demonstrated the striking capability of the SCAP system in obtaining highly purified and enriched SPIN1/SPINDOC complexes and use of the extended crosslinking analysis pipeline, SCAP-XL, in integrative structural modeling. The SCAP method, in principle, could be also applied to dissect subpopulations for proteins with more complicated interactions by enriching for co-associations of both bait proteins.

In this study, we demonstrate another application of SCAP technology using Halo-WDR76 and SNAP-SPIN1 as bait proteins (**Fig. 1C**). Here, we first co-expressed bait proteins in cells depleted of endogenous WDR76. Using the modified SCAP-MS and SCAP-XL pipelines, we separated the WDR76/SPIN1 complex from other WDR76 complexes containing CBX proteins and the SPIN1/SPINDOC complexes; identified co-associated proteins; and also identified direct physical interactions between the protein complex and various histones. In contrast to facing hundreds of possible interactions in conventional AP-MS analysis, the SCAP technology allowed us to focus on 27 most enriched co-associated proteins of WDR76 and SPIN1, providing more precise inference of WDR76/SPIN1 function in DDR for future studies.

## Results

### Results of WDR76 and SPIN1 Single-Bait AP-MS

We first analyzed interactions of WDR76 and SPIN1 separately using Halo-tag AP-MS. Our updated WDR76 AP-MS results are mostly consistent with previous data (*9, 10*). In this experiment, we identified 395 proteins significantly enriched over control purifications (**Data S2A**). Among the top 20 proteins ranked by abundance we detected histones (H2A, H2B, H3, H4), CBX proteins (CBX1, CBX3, CBX5) and SPIN1 (**Fig. 1B**). GAN (gigaxonin) appeared the 21^st^ in the list. The chaperonin containing TCP1 complex (CCT or TriC complex) members were the next abundant proteins enriched in the list. A total of eight components of CCT complex subunits (TCP1, CCT2, CCT4, CCT5, CCT7, CCT3, CCT8, CCT6A) were ranked within top 50 by dNSAF values. We also detected other previously reported interactions, such as HELLS, SIRT1, HDAC1, HDAC2, and proteins involved in DNA damage repair (XRCC5, XRCC6, PARP1, PRKDC, DDB1, RAD23B) (*9, 10*). In agreement with a published data which suggested a role of WDR76 in targeting proteins for ubiquitination and degradation (*18–22*), we discovered several ubiquitination related proteins including DCAF7, USP7, USP11, USP9X, HUWE1 and UBR5. Taken together, these data alongside previous reports suggest the diverse protein interaction networks associated with WDR76.

In our new AP-MS data of SPIN1, we discovered more co-purified proteins than prior experiments (*1*). A total of 292 proteins were significantly enriched (**Data S2A**). SPINDOC is the most abundant protein other than the bait protein, SPIN1 (**Fig. 1B**). WDR76 was also amongst the most abundant proteins with an average dNSAF value of 0.017. Other interesting proteins in the top 20 most abundant proteins included histone H3, lamin-B receptor (LBR) that links chromatin to nuclear envelope (*35*), and several mRNA processing and export related proteins (ALYREF, THRAP3, BCLAF1, ERH) (**Fig. 1B**). In addition, proteins involved in histone methyltransferase complexes (KMT2A, KMT2B, RBBP5, DPY30, ASH2L, PRMT1, FBL, MEN1) and cell cycle related proteins (MCM proteins and RPA proteins) copurified with Halo-SPIN1(**Data S2A**). However, no functional implication of SPIN1 and WDR76 together became apparent by comparing the single-bait AP-MS results of two bait proteins. Thus, the SCAP method might be necessary to gain more insights on the biological meaning of a subset of interactions.

### The *in vivo* Interaction between WDR76 and SPIN1

Halo tag (*36*) and SNAP tag (*37*) have been selected as the affinity tags in SCAP, which allows for convenient differential imaging of the bait proteins. Before applying SCAP to purify WDR76 and SPIN1, we took advantage of the imaging feature to validate the interaction between Halo-WDR76 and SNAP-SPIN1 in live cells using acceptor photobleaching Förster resonance energy transfer (apFRET) assay (*38*) and fluorescence cross-correlation spectroscopy (FCCS) assay (*39*). Similar to assessing SPIN1 and SPINDOC by Liu *et al.* (*1*), Halo-WDR76 and SNAP-SPIN1 were co-expressed, stained with the corresponding fluorescent ligands and imaged in live cells (**Fig. 2**). In the apFRET assay, we observed about 2% of FRET efficiency between Halo-WDR76 and SNAP-SPIN1, which was significantly higher than that measured with SNAP-Control (**Fig. 2B**). In the FCCS assay, we observed a cross correlation of Halo-WDR76 and SNAP-SPIN1, but not Halo-WDR76 and SNAP-Control (**Fig. 2D**). The fraction of Halo-WDR76 binding to SNAP-SPIN1 was also calculated using y amplitudes of self- and cross-correlation curves (**Fig. 2E**). In summary, the positive FRET result indicated a direct interaction of WDR76 and SPIN1 *in vivo* and the FCCS result suggested WDR76 and SPIN1 co-diffuse in a complex. The FCCS result was particularly intriguing because it also suggested that the complex formed by WDR76 and SPIN1 is mobile in the nucleus. As we mentioned above, both WDR76 and SPIN1 may interact with core histones (**Fig. 1B and Data S2A**). Moreover, chromatin immunoprecipitation (ChIP) assays have indicated that both WDR76 (*17*) and SPIN1 (*28*) bind to chromatin. Since chromatin associated species are usually less mobile and difficult to measure by FCCS, our result indicated that WDR76 and SPIN1 might form a complex independent of their binding to chromatin. This result also raised the question whether WDR76/SPIN1 in complex would bind to histones although histone H3 was enriched in both WDR76 and SPIN1 purifications (**Fig. 1B**).

**Fig. 2.**
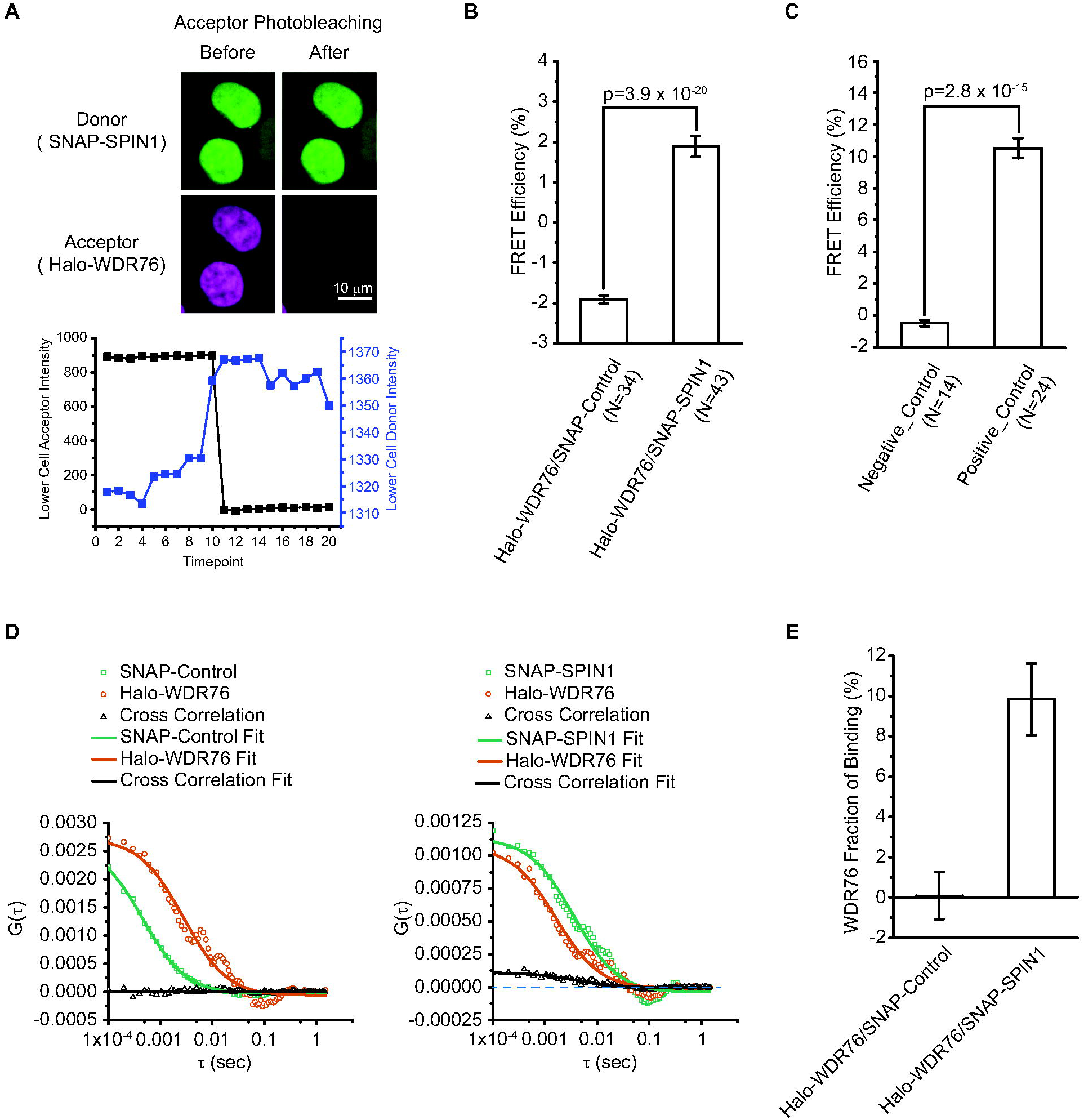
Imaging approaches to assess the interaction between WDR76 and SPIN1. The interaction between WDR76 and SPIN1 was characterized using Acceptor-Photobleaching Fluorescence Resonance Energy Transfer (apFRET) (**A-C**) and Fluorescence Cross-Correlation Spectroscopy (FCCS) (**D-E**). (**A**) Example image and intensity measurement of apFRET. Halo-WDR76 was labeled with HaloTag TMRDirect ligands and SNAP-SPIN1 was labeled with SNAP-Cell 505-Star ligands. (**B**) Averaged FRET efficiencies measured for Halo-WDR76 and SNAP-SPIN1 in live HEK293FRT cells. SNAP tag was used as donor in the control experiment to be tested with Halo-WDR76. Error bars stand for standard error of mean and p-values were calculated using two-tailed t-test. (**C**) FRET efficiencies measured for control proteins in live HEK293FRT cells. Separate Halo tag and SNAP tag were used as a negative control pair and a fusion protein of Halo tag and SNAP tag was used as positive control. Error bars stand for standard error of mean and p-values were calculated using two-tailed t-test. (**D**) Auto and cross-correlation curves for Halo-WDR76 and SNAP-SPIN1, and for Halo-WDR76 and SNAP tag as control experiment. (**E**) The average percentage of Halo-WDR76 binding to SNAP tag or SNAP-SPIN1 was calculated from the y amplitudes of correlation curves. Error bars stand for standard error of mean. Values was obtained based on the fitted correlation curves, t-test was not performed for this data.

### SCAP Analysis of WDR76 and SPIN1

In the formerly described SCAP workflow, we generated cell lines stably expressing both bait proteins using a dual-expression plasmid from wildtype parental cells, in which the endogenous bait proteins remain present (*1*). Using the stable-expressing cells, proteins from whole cell extracts were first isolated on SNAP affinity beads and then eluted using PreScission protease to obtain fraction E1. Then E1 was purified in tandem using Halo affinity beads. The unbound supernatant of the Halo purification was collected as fraction UB2. The proteins captured by the Halo beads were eluted using the TEV protease as fraction E2. One major concern of using such cell lines to perform SCAP is the interfere from the endogenous version of the bait protein for the later purification step. These untagged proteins might be captured with the SNAP-tagged bait, but cannot be captured during Halo purification, thus a portion of co-associated proteins would be lost in the final fraction E2. Though the original SCAP protocol still functioned for SPIN1 and SPINDOC (*1*), removing untagged version of the Halo bait protein from the system should theoretically enhance the yield of SCAP. For WDR76 and SPIN1, we first depleted the endogenous WDR76 (the Halo-tagged bait for the second purification in SCAP) from HEK293 cells bearing a FRT recombination site by deleting both alleles of the corresponding gene, and then we built cell line expressing Halo-WDR76 and SNAP-SPIN1 in WDR76 knockout background (**Fig. 1C**). Using these cells, the WDR76 captured in E1 were completely tagged by Halo and enriched by serial capture. In addition, we optimized several steps of the SCAP protocol: we tripled the number of cells to extract proteins at a higher yield; we subjected 90% of E1 to subsequent purification to use more material in the final enrichment; we also adjusted conditions and timing of purification steps. Comparable amount of E1, UB2 and E2 samples from each SCAP experiment were analyzed with label-free quantitative proteomics (SCAP-MS), multidimensional protein identification technology (MudPIT) (*40*) (**Data S2B** and **Fig. S1**). The modified SCAP-MS workflow is demonstrated in **Fig. 3A**. For a better comparison of SCAP-MS and single-bait AP-MS of WDR76, we also generated a stable cell line that expresses Halo-WDR76 alone in the same WDR76 knockout background to perform AP-MS (**Data S2C**).

**Fig. 3.**
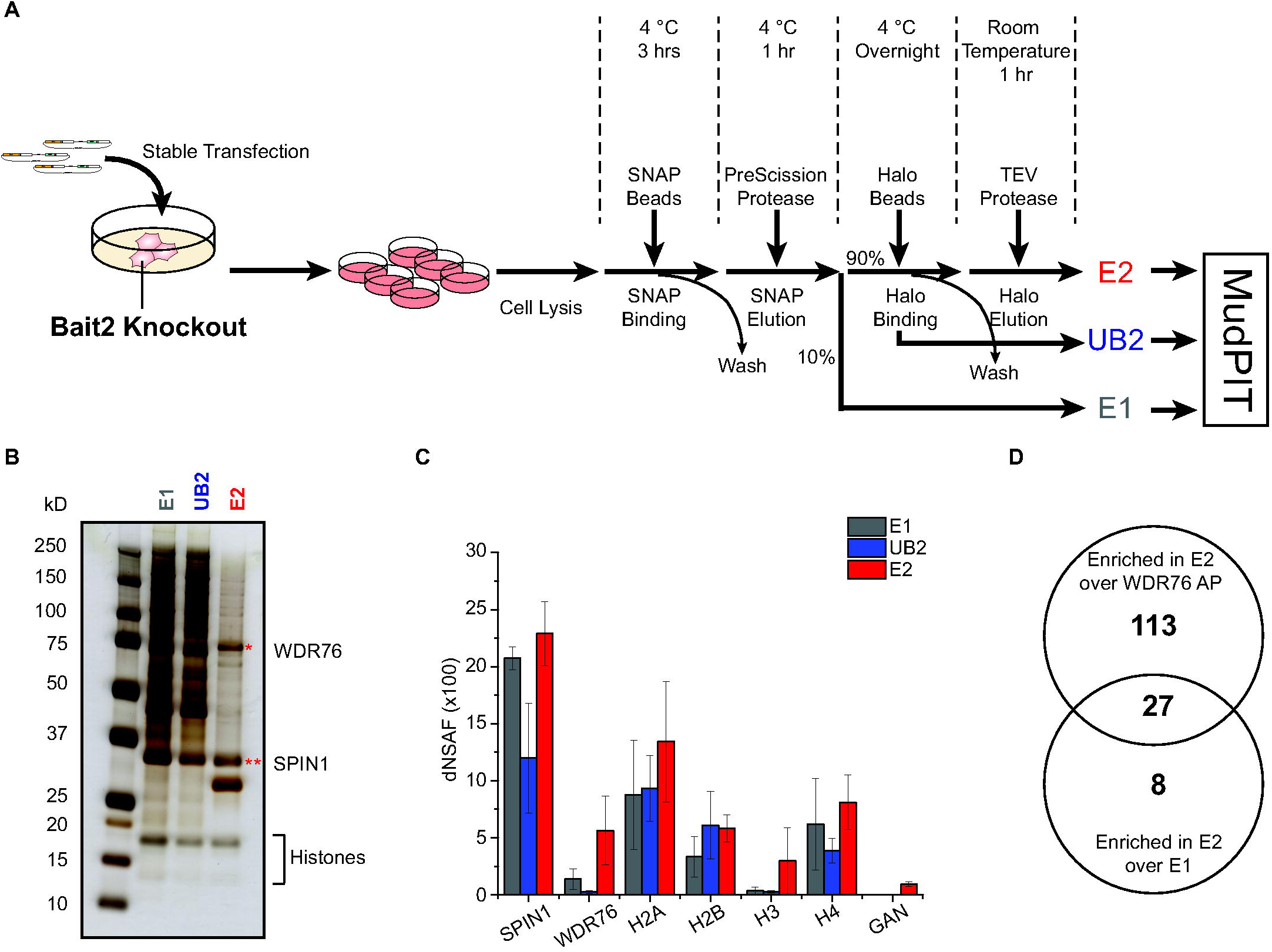
Analyzing co-purified proteins of WDR76 and SPIN1 with a modified SCAP workflow followed by Multidimensional Protein Identification Technology (MudPIT). (**A**) A modified workflow of Serial Capture Affinity Purification followed by quantitative proteomics analysis (SCAP-MS). For obtaining materials for SCAP, stable transfection of the dual-expression vector is performed on cells deleted of the endogenous bait protein in the second step of purification (Bait2). SCAP purification is similar to the original workflow (*1*) with minor optimization. E1, UB2 and E2 fractions were all separately analyzed by MudPIT and dNSAF values were calculated from spectra counts for all identified proteins. (**B**) Silver stained SDS-PAGE of the proteins in the E1, UB2, and E2 fractions of SCAP using Halo-WDR76 and SNAP-SPIN1 as bait proteins. The anticipated band of WDR76 is marked with * and SPIN1 is marked with ** based on their sizes. Bands of histones are expected to appear in the range below 20kD. (**C**) The dNSAF values of WDR76, SPIN1, GAN and histones in E1, UB2, and E2 fractions. Each dNSAF value plotted is the average from three biological replicates; error bars represent standard deviations. (**D**) Venn diagram showing numbers of proteins enriched in E2 fraction over E1 and enriched by SCAP over single-bait WDR76 purification. There are 27 proteins shared by both groups of enriched proteins.

The modified SCAP method again showed the striking capability to reduce sample complexity (**Fig. 3B**), illustrated by a much cleaner E2 sample comparing to E1. In this SCAP-MS analysis of WDR76 and SPIN1, we detected in total 699 proteins in at least one of E1 or E2 fraction and 165 proteins in E2 were significantly enriched over the blank control purification (**Data S2B & F**). The top 20 proteins ranked by dNSAF values in E2 are shown in **Fig. S1C**. Besides the high abundance of both bait proteins in E2, other proteins including histones and GAN were also among the top 20 identifications, suggesting their strong association with both WDR76 and SPIN1. Upon comparing the three fractions, WDR76, GAN and histones appeared more abundant in E2 than E1 (**Fig. 3C**). By comparing E1 and E2 statistically using QSPEC (*41*), WDR76, GAN, histone H2A, H2B and H3 were significantly enriched in E2 over E1 (**Data S2B & F**). The other bait SPIN1 was not further enriched during the 2^nd^ purification step (**Fig. 3C**), which could be explained by that only a fraction of SPIN1 was bound by WDR76 in cells. In addition to the most abundant detections, we also identified other co-captured proteins of WDR76 and SPIN1 in E2, which might also provide clues for the functions of the complex. Among the 165 proteins enriched in E2 over control, 35 proteins were significantly enriched in E2 over E1 and 140 over single-bait Halo-WDR76 co-purified samples (**Data S2C and F**). Binary analysis of the 35 and 140 proteins found 27 proteins of overlap, which were enriched in E2 over both E1 and WDR76 purification (**Fig. 3D** and **Data S2F**). As expected, the CBX proteins were no longer significantly enriched by SCAP comparing to Halo-WDR76 purification along, demonstrating that the SCAP method is capable of distinguishing a subpopulation of WDR76 that are in complex with SPIN1.

The major components in E2, including SPIN1, WDR76, all four canonical histones (H2A, H2B, H3, H4) and GAN, constituted about 60% of the total sample (summed average dNSAF value: 0.598). This enrichment was similar in the SCAP-MS analysis of SPIN1 and SPINDOC, in which SPIN1/SPINDOC complex was also about 60% of total proteins in E2 (*1*). Intriguingly, taken together the SCAP-MS results of Halo-WDR76/SNAP-SPIN1 and Halo-SPIN1/SNAP-SPINDOC (*1*) also suggested that SCAP method successfully distinguished two SPIN1-containing protein complexes (**Data S2D**). When comparing to the single-bait Halo-SPIN1 purification, WDR76 was significantly enriched by SCAP using both Halo-WDR76 and SNAP-SPIN1 as baits, indicating the successful purification of WDR76/SPIN1 over other SPIN1 interactions. In contrast, the SPINDOC co-purified with SNAP-SPIN1 in E1 was barely captured during the second step of WDR76/SPIN1 SCAP (dNSAF in E2: 0.005) comparing to unbound fraction UB2 (**Fig. S1**). In the SPIN1/SPINDOC SCAP-MS analysis we published previously (*1*), WDR76 was not detected in E2 at all (**Data S2D**). These results suggested the presence of WDR76 and SPINDOC to be exclusive when forming complexes with SPIN1.

SPIN1 binds H3K4me3 (*23, 25, 26*) and various histones were identified in WDR76 AP-MS analyses (*8–10*). Besides published results, the single-bait AP-MS data in this study showed that WDR76 co-purified with all 4 types of core histones while SPIN1 mostly co-purified histone H3 (**Fig. 4A** and **Data S2A-C**). In the SCAP-MS analysis of WDR76/SPIN1, core histones (H2A, H2B, H3 and H4) were also among the most abundant co-captured proteins of WDR76 and SPIN1 (**Fig. 3C** and **Data S2B**). In addition, Histone H2A, H2B and H3 were in the 27 proteins more enriched in E2 over both E1 and WDR76 purification (**Data S2F**). These results suggested that WDR76 and SPIN1, when forming a complex, were still able to interact with histones. Since the complex co-purified with all 4 types of histones as WDR76 but SPIN1 alone captured less histone H2A, H2B and H4 (**Fig. 4A**), the binding of SPIN1 to WDR76 did not deprive the interactions of WDR76 with histones. Furthermore, as we noted other WDR76 co-purified proteins, the CBX proteins (**Fig. 1B**), recognize opposite post translational modifications (PTMs) on histone H3 from SPIN1 (*30–32*), we raised possibility that WDR76 might interact with differently modified histones when associated with different partners. With the AP-MS and SCAP-MS data above, we were able to analyze the PTMs on H3K9 (**Data S2E**). As shown in **Fig. 4B Error! Reference source not found.**, high levels of acetylation on H3K9, which is also considered transcriptionally active modifications that sometimes co-exist with H3K4me3, were detected in the E2 fraction of WDR76/SPIN1 SCAP, but not in single-bait WDR76 purifications. While trimethylation levels on H3K4 could not be measured due to this short peptide being difficult to identify by LC/MS, we tested the presence of H3K4me3 in WDR76/SPIN1 co-purified proteins using SCAP followed by Western blot analysis (**Fig. 4C**). In comparison, we analyzed SPIN1/SPINDOC SCAP along with WDR76/SPIN1. As shown in **Fig. 4C**, H3K4me4 was clearly co-purified with WDR76/SPIN1 but was absent for SPIN1/SPINDOC. This observation is consistent with our previous conclusion that the binding of SPINDOC to SPIN1 could interrupt the binding of SPIN1 to H3K4me3 (*1*). Taken together, WDR76/SPIN1 in complex selectively bound with histone H3 bearing specific PTMs.

**Fig. 4.**
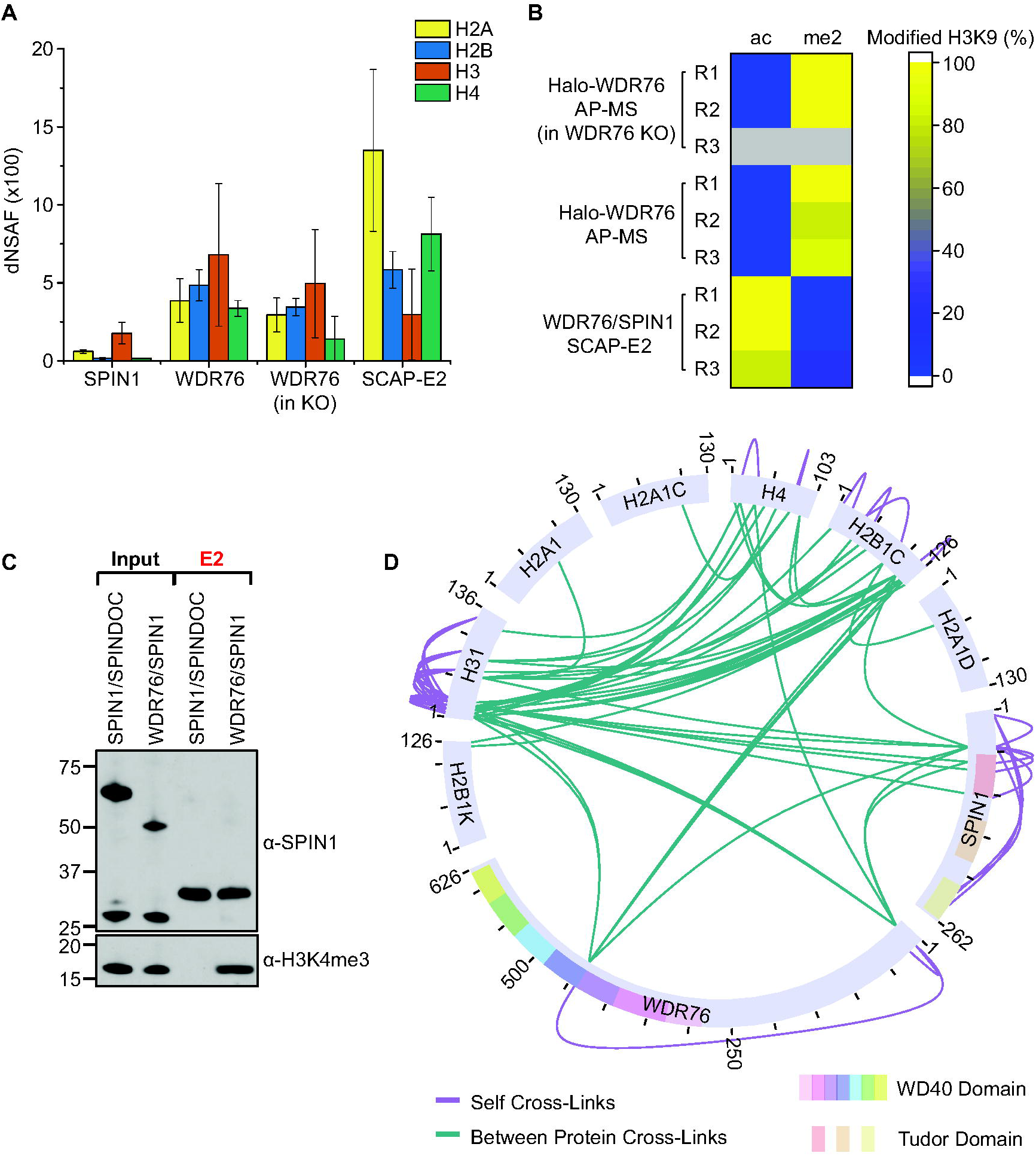
Charactering the interactions between the WDR76/SPIN1 protein complex and different types of core histones with SCAP-MS and SCAP-XL. (**A**) The dNSAF values of core histones in samples obtained from different purifications. All single-bait purifications were Halo purifications. Halo-WDR76 purification was performed either with or without the present of endogenous WDR76. SNAP-SPIN1 and Halo-WDR76 were baits for SCAP. Each dNSAF value plotted is the average from three biological replicates; error bars represent standard deviations. (**B**) Distinct post-translational modification status of H3K9 on histone H3 captured in SCAP E2 fraction versus that of histone H3 captured in single-bait purification using WDR76 as bait. The percentages of modified H3K9 were calculated from spectral counts. Three replicates for each type of experiment are shown. (**C**) Comparison of lysine 4 trimethylated histone H3 co-purified with WDR76/SPIN1 or SPIN1/SPINDOC complex by SCAP. The shared bait protein, SPIN1, was detected by antibodies recognize both endogenous and tagged versions of SPIN1. SCAP purification was performed from extracts of 293FRT cells expressing SNAP-Flag-SPIN1 + Halo-HA-WDR76 and Halo-HA-SPIN1 + SNAP-Flag-SPINDOC. Input and eluted fractions were resolved on gels followed by silver staining and western blotting. The expected band for the target proteins are depicted by an asterix. The present of H3K4me3 was detected by antibodies specifically recognize histones carrying the modification. This result was validated by two other repeated sets of purifications. (**D**) Direct interactions between WDR76, SPIN1 and various types of histones revealed by SCAP coupled with crosslinking mass spectrometry analysis (SCAP-XL). The crosslinks were visualized in circle view generated with xiView (*42*).

### A SPIN1/WDR76 Complex that Directly Interacting with Histones Distinguished from SPIN1/ SPINDOC

To identify direct interactions and to gain structural insights on the WDR76/SPIN1 complex, we next implemented SCAP-XL, where we first performed the modified SCAP for Halo-WDR76 and SNAP-SPIN1 followed by crosslinking reaction on samples using a MS-cleavable crosslinker, DSSO (disuccinimidyl sulfoxide) (**Fig. S2A**). Then we analyzed crosslinked E2 fraction by crosslinking mass spectrometry (XL-MS), and visualized the data using xiVIEW (*42*) (**Data S3A**). Detected crosslinks to WDR76, SPIN1 and histones are shown in circular view (**Fig. 4D**). WDR76 and SPIN1 were crosslinked to each other, which confirmed their direct interaction as we also proved by an apFRET assay (**Fig. 2B**). Histone H2B, H3 and H4 from the list of top identifications in SCAP-MS were also crosslinked to one or both baits, suggesting that WDR76 and SPIN1 directly interact with those types of histones. Histone H2B and H3 were found to crosslinked to both WDR76 and SPIN1 with multiple crosslinks, while H4 was only crosslinked to WDR76 with one crosslink. Histone H2A, despite of the high abundance in co-captured proteins of WDR76 and SPIN1 (**Fig. 3C**), was not found directly crosslinked to either bait. However, a number of crosslinks were detected between histones, thus H2A might interact with the WDR76/SPIN1 complex indirectly through other histones. Histone H3 was crosslinked to WDR76 and SPIN1 with its N terminal region, in contrast, H2B was crosslinked to the baits with its C terminal region. Crosslinking sites of SPIN1 distributed mostly near the N terminal region, while no crosslinking site fell in the second Tudor domain, where the pocket to bind H3K4me3 locates (*25, 26*). In contrast to the SCAP-XL result, the XL-MS analysis of Halo-WDR76 purification did not identify crosslinked peptides from SPIN1 (**Fig. S2C** and **Data S3D**). Histone H2B, H3 and H4 were crosslinked to WDR76, however, only H3 was detected with more than one crosslinks (**Fig. S2C**).

Next, we performed structural modeling for tridimensional visualization of crosslinks. Although several crystal structures of SPIN1 have been published (*24–26*), the N terminal region, where some critical crosslinked lysine fell in, was not covered by these structures. Since the structures of both full-length WDR76 and SPIN1 are not available, we performed *ab initio* prediction of their structures with I-Tasser (*43, 44*) from amino sequences. I-Tasser provided 5 models for each protein. Intra-crosslinks were mapped to the predicted models (**Fig. S3A & B**) and distances between crosslinked residues were measured (**Data S4A**). For WDR76, we obtained two proper folded models, model3 and model5, and 5 out of 6 detected intra-crosslinks can be satisfied in both models (**Fig. S3A**). For SPIN1, models are more similar to each other (**Fig. S3B**). The model2 matches all intra-SPIN1 crosslinks (**Data S4A**), therefore was selected as our best model for docking with WDR76. This model can also be largely aligned to a known SPIN1 structure in PDB: 4mzf (*26*) (**Fig. S3C**). After filtering inter-crosslinks between WDR76 and SPIN1 with DisVis (*45, 46*) (**Data S4B**), we chose WDR76 model3 to dock with SPIN1 model2. The docking was performed with HADDOCK 2.4 webserver (*47, 48*) using the 3 inter-crosslinks between WDR76 and SPIN1 as restraints (**Data S4C**). The best four models from the top cluster with the lowest HADDOCK score are shown in ensembled representation (**Fig. S4A**). Crosslinks were mapped to these models and the distance restraints were all satisfied by 4 models (**Data S4D**). The model with the best energy was used as an example for visualization (**Fig. S4B & C**). From **Fig. S4B**, we noticed that the interface between WDR76 and SPIN1 in this model did not overlap with the Tudor domains of SPIN1. Then we aligned the WDR76/SPIN1 model to a known SPIN1 structure containing a methylated histone H3 peptide (PDB: 4mzf) (*26*) in **Fig. S4C** to see whether WDR76 would interfere with the binding of SPIN1 to H3. The histone H3K4me3R8me2a binding pockets of SPIN1 indicated by Su *et al.* (*26*) was not covered by WDR76 in the complex model (**Fig. 5A**). In contrast, this binding pocket is covered in the SPIN1/SPINDOC complex model (**Fig. 5B**).

**Fig. 5.**
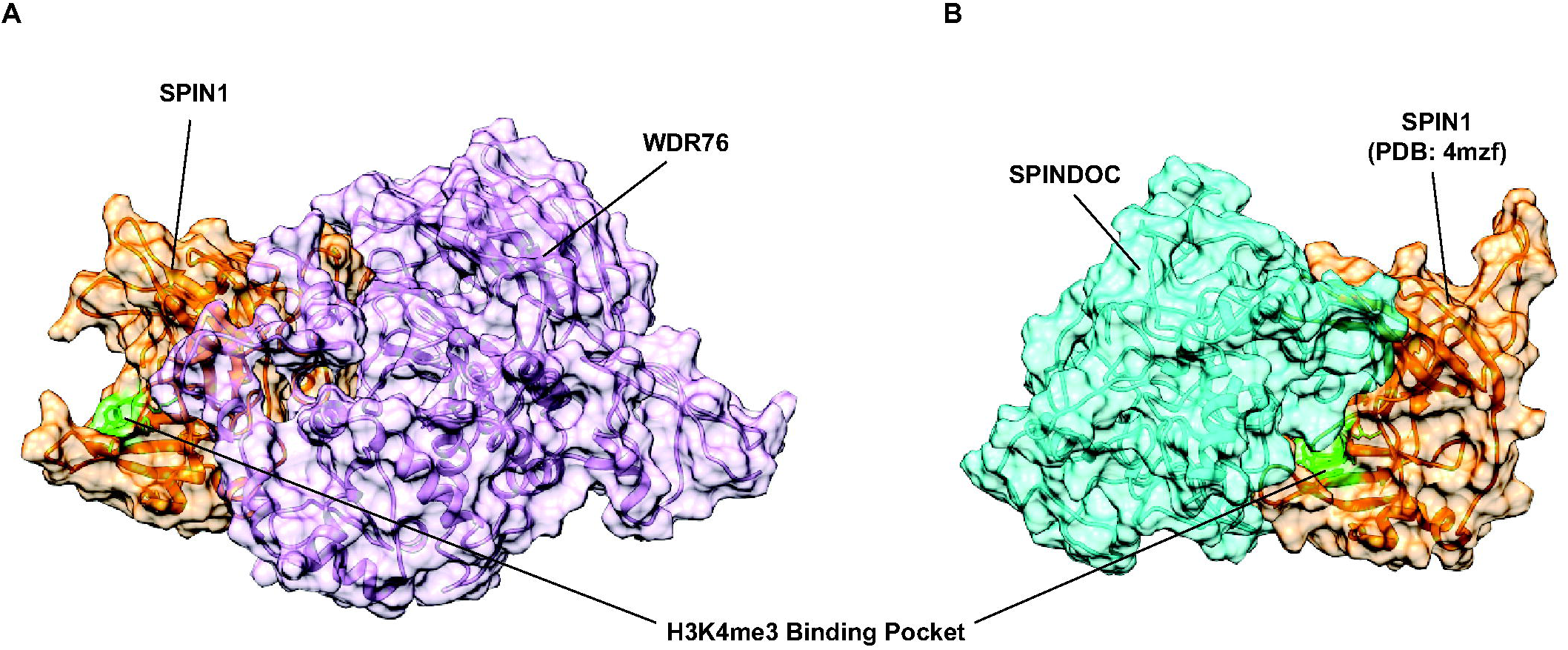
Comparing the structural models of two distinct SPIN1 containing complexes. (**A**) The heterodimer model of WDR76 and SPIN1 with the lowest HADDOCK score is displayed in surface view. WDR76 is shown in plum color and SPIN1 is shown in orange. (**B**) The best heterodimer model of SPIN1 and SPINDOC described by Liu *et al.* (*1*) is displayed in surface view. SPINDOC is shown in cyan. SPIN1 chain in this model was adopted from PDB: 4mzf and is also colored in orange. In both complex models, the H3K4me3 binding of SPIN1 are marked in green.

### Function of WDR76/SPIN1 in the DNA Damage Response

Besides histones, SCAP-MS analysis of WDR76 and SPIN1 identified other less abundant co-captured proteins enriched in E2. Noticeably, both XRCC5 and XRCC6 were among the 27 proteins more enriched in E2 over both E1 and WDR76 single-bait purification (**Data S2E**). XRCC5 and XRCC6 forms heterodimer (Ku complex) that binds double-strand DNA. A well-known function of the XRCC5/XRCC6 dimer is that it is recruited to DNA double-strand breaks (DSB) in the Non-Homologous DNA End Joining (NHEJ) pathway (*49*). Other proteins involved in the NHEJ pathway, DNA-PKcs (PRKDC) and PARP1, were also identified by SCAP-MS (**Fig. 6A**). This result strongly implied a potential function of the WDR76/SPIN1 complex in NHEJ. WDR76 has been reported to be involved in DNA damage response (DDR) and factors in NHEJ repair pathway were identified by WDR76 AP-MS (*9–14*). SPIN1, however, has never been linked to DDR in existing data, besides its interaction with WDR76. Our data showed that XRCC5/XRCC6 were co-associated proteins of both WDR76 and SPIN1, suggesting a potential role of SPIN1 in DDR as well.

**Fig. 6.**
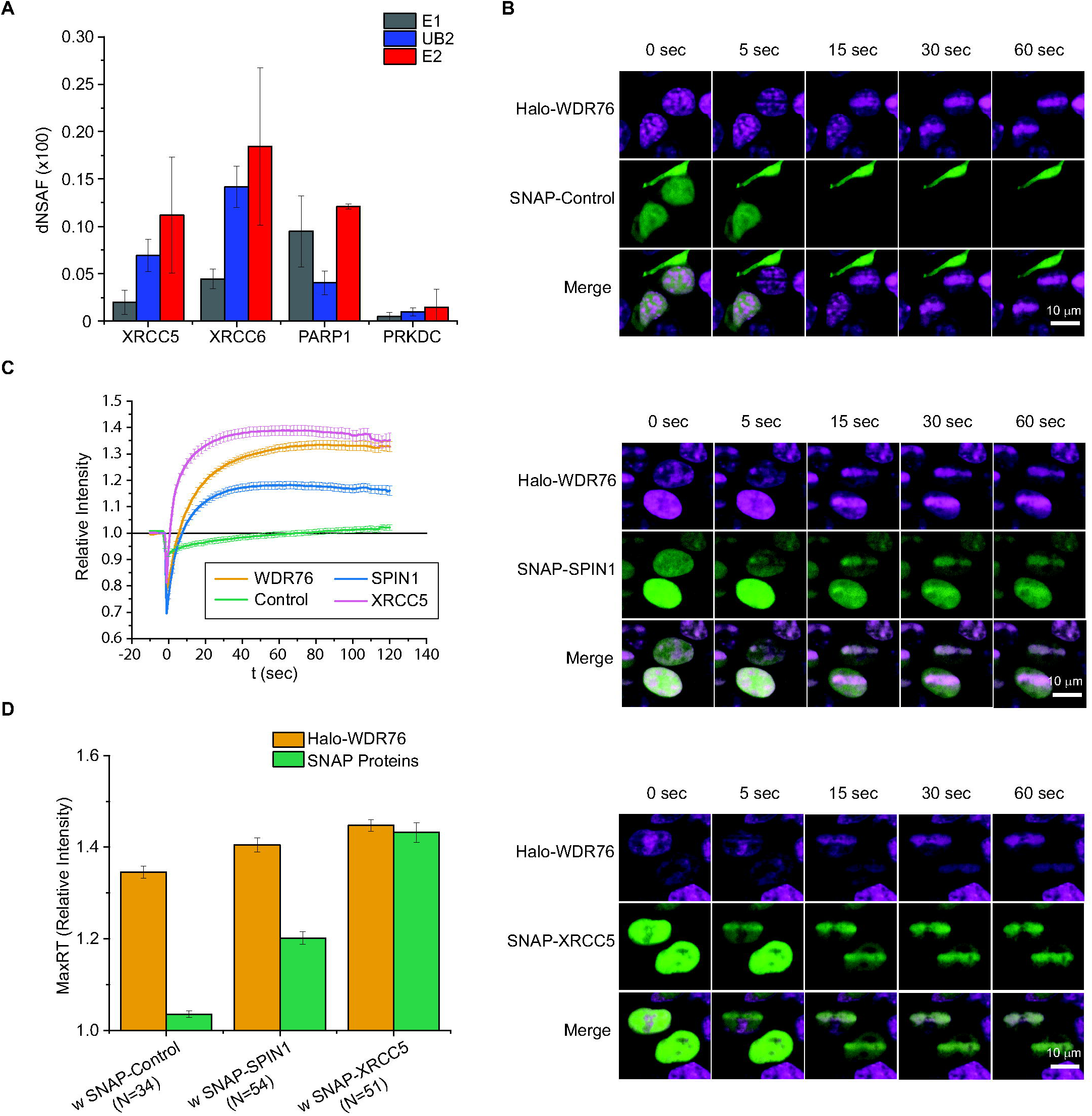
Potential function of WDR76/SPIN1 complex in DNA Damage Response (DDR). (**A**) Identification of proteins involved in non-homologous DNA end joining (NHEJ). Each dNSAF value plotted is the average from three biological replicates; error bars represent standard deviations. (**B**) Examples of time course imaging. In each experiment, a SNAP-tagged protein was co-expressed with Halo-WDR76. Halo-tagged proteins were labeled with HaloTag TMRDirect ligands and SNAP-tagged proteins were labeled with SNAP-Cell 505-Star ligands. Cells were also stained with HOECHST 33258. 405nm UV laser damage was introduced at timepoint 0 in a stripe-shaped confined area. (**C**) Recruitment of Halo- or SNAP-tagged proteins to laser induced damages. Recruitment level is presented by relative intensity at the stripe, calculated from the intensity of each timepoint normalized to the intensity before microirradiation. For Halo-WDR76, the sample co-expressed with SNAP-Control is plotted. Each data point is an average from all measured cells; error bars stand for standard error of mean. (**D**) The maximum recruitment of each protein within 2min after microirradiation. Each maximum recruitment value is an average from all measured cells; error bars stand for standard error of mean.

To test whether SPIN1 also responds to DNA damage in live cells, we applied an imaging-based recruitment assay (*9*). In this assay, Halo- or SNAP-tagged proteins were expressed in HEK293 cells and labeled with the corresponding fluorescent ligands. Cells were also treated with Hoechst33258 as sensitizer. Then we performed microirradiation treatment, in which we induced DNA damage in a defined area on each cell using a 405nm laser. After microirradiation, the treated cell was recorded for up to 2 minutes (**Fig. 6B**). Using this assay, we have previously reported the recruitment of WDR76 and XRCC5 to the irradiated region (*9*). Consistent with these previous findings, we now observed that WDR76 and XRCC5 were recruited to the damaged region within 30 seconds (**Fig. 6C**). Similarly, SNAP-tagged SPIN1 was also rapidly enriched in the laser damaged region (**Fig. 6C & D**). Although the detailed mechanisms remained unclear, this result supports the hypothesis that both WDR76 and SPIN1 are involved in DDR. Intriguingly, the recruitment of SPIN1 was nearly concurrent with WDR76 but obviously slower than XRCC5 (**Fig. 6C**). To compare the kinetics of recruitments, we performed nonlinear fitting for the average Rt values of each protein and estimated the rate of recruitment using the time taken to reach half of MaxRT (t_MaxRT/2_) (**Fig. S5** and **Data S5E**). The measurements showed that there was only 0.7 second of difference between the t_MaxRT/2_ of WDR76 and SPIN1, versus XRCC5 was about 4 second quicker than WDR76 to reach the half of its MaxRT. This result suggested that SPIN1 and WDR76 might be recruited to DNA damage site as a complex through interaction with the Ku complex. This is also consistent with our FCCS result, which suggested that WDR76 and SPIN1 in complex were mobile in the nucleus.

## Discussion

Here, we described a modified Serial Capture Affinity Purification (SCAP) system (**Fig. 1C**), in which we first deleted the endogenous version of bait protein used in the 2^nd^ purification step, to improve the yield of the serial capture. Using the modified SCAP method, we analyzed WDR76 and SPIN1 by live cell imaging, quantitative proteomics and crosslinking mass spectrometry. Results were summarized in **Fig. 7A**. We reported that the core histones and GAN were among the top abundant co-captured proteins of WDR76 and SPIN1 in the SCAP E2 fraction. We confirmed the direct interaction between WDR76 and SPIN1 (**Fig. 7B**). Histones H2B, H3 and H4 were also found to interact directly with SPIN1 or WDR76 (**Fig. 7B**). Finally, XRCC5 and XRCC6 were enriched by WDR76/SPIN1 SCAP, suggesting a role of the WDR76/SPIN1-containing complex in DNA damage response.

**Fig. 7.**
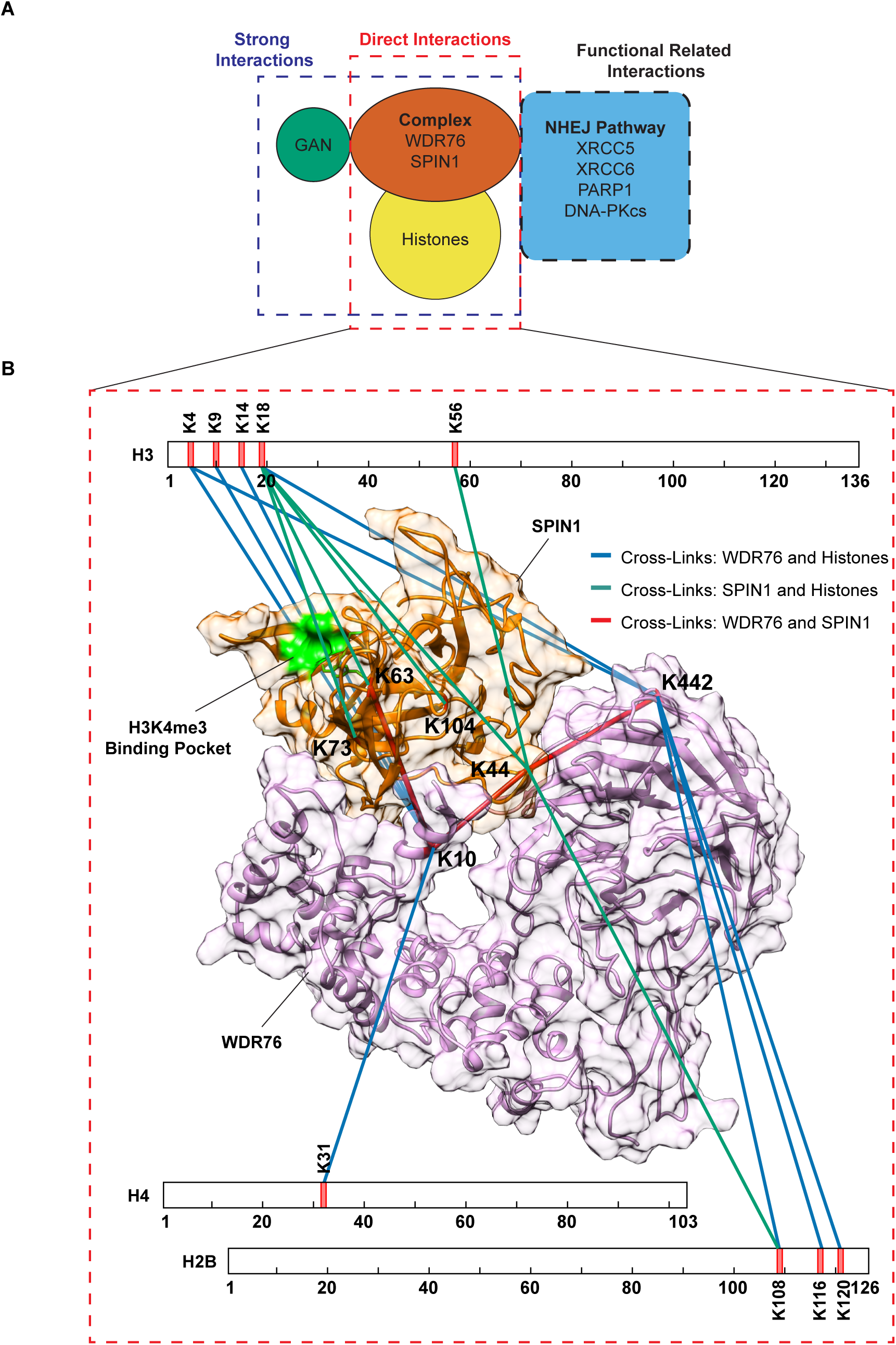
Model of SCAP-MS and SCAP-XL results using Halo-WDR76 and SNAP-SPIN1 as bait proteins. (**A**) From the quantitative analysis of SCAP samples, GAN and core histones were identified as strong interactions of WDR76/SPIN1 complex based on their abundance in E2 fraction. Low abundant interactors, including XRCC5, XRCC6, DNA-PKcs and PARP1, indicated the potential roles of both WDR76 and SPIN1 in DNA damage response. DSSO crosslinking of SCAP purified proteins followed by mass spectrometry analysis further supported the direct physical interaction between WDR76 and SPIN1. Crosslinks between WDR76/SPIN1 and histones also suggested direct protein-protein interactions. (**B**) Visualization of direct crosslinks to WDR76 or SPIN1. WDR76/SPIN1 complex is displayed by a predicted structural model in surface view, in which WDR76 is colored in plum color and SPIN1 is colored in orange. The H3K4me3 binding pocket of SPIN1 is colored in green. All crosslinked lysine residuals were labeled and marked in red. Crosslinks between WDR76 and SPIN1 are shown as red lines. Crosslinks between histones and WDR76 are shown as blue lines, while those between histones and SPIN1 are shown as green lines.

WDR76 was first identified in yeast through AP-MS analysis of core histones (*8*). Its amino acid sequence contains seven WD40 domain repeats and is conserved in higher eukaryotes (*10*). Comprehensive AP-MS studies of human WDR76 identified many legitimate interactions (*9, 10*), yet its function remains enigmatic. One reason is that WDR76 might conduct various activities with different binding partners (**Fig. 1A**). This multiplicity is difficult to resolve by the conventional AP-MS approach, but is a good test case to demonstrate the power of SCAP technology. Among the WDR76 co-purified proteins, we chose SPIN1 as the other bait protein to perform SCAP. SPIN1 binds H3K4me3 with its second Tudor domain (*25, 26*), thus its interaction with WDR76 inferred their collaborative function in chromatin related activities. In consistent with existing knowledge, core histones appeared as major co-associated proteins of WDR76 and SPIN1 enriched by SCAP (**Fig. 3C**). In addition, SCAP successfully enriched 35 co-captured proteins of both WDR76 and SPIN1 in E2 fraction from E1. By further contrasting against single-bait WDR76 AP-MS, 27 proteins were found more enriched by SCAP. The list of candidate co-interactors was greatly shortened (**Fig. 3D**). XRCC5 and XRCC6 were easily noticed from only 27 proteins, implying the potential function of both WDR76 and SPIN1 in DDR. We performed a straightforward recruitment assay, where SPIN1 was for the first time tested to response upon DNA damage (**Fig. 6B-D**).

SCAP-XL of WDR76 and SPIN1 provided evidence for direct protein-protein interactions (**Fig. 4D**). No crosslinking site was observed in the second Tudor domain of SPIN1 in this analysis (**Fig. 4D**), indicating that the interaction of SPIN1 with WDR76 might not prevent it from binding to H3K4me3. This is consistent with the conclusion from PTM analysis of histone H3 by SCAP coupled with mass spectrometry or Western blot (**Fig. 4B and C**), that the WDR76/SPIN1 complex might selectively bind certain epigenetic markers through SPIN1. As extra supporting evidence, the H3K4me3 binding pocket of SPIN1 has no overlap with the interface with WDR76 in the model (**Fig. 5B**). Moreover, WDR76 was crosslinked to H3K4 and H3K9, which are likely to be modified in SPIN1-bound histone H3. This interesting observation suggested the possibility for WDR76 and SPIN1 in the complex binding two different copies of histone H3.

Another SPIN1-containing complex, the SPIN1/SPINDOC complex, was previously analyzed using SCAP technology (*1*). The SCAP-MS results suggested that the complex SPIN1 formed with WDR76 is a distinct complex from the one formed with SPINDOC (**Data S2D**). In contrast to the WDR76/SPIN1 complex, SPIN1 lost binding of histone H3 in the SPIN1/SPINDOC complex. This is also supported by structural modeling results: in the SPIN1/SPINDOC complex, SPINDOC blocks the H3K4me3 binding pocket of SPIN1; versus the pocket is left open in the WDR76/SPIN1 model (**Fig. 5**). It worth noting that in this study we showed evidence of WDR76/SPIN1 being recruited to laser induced DNA damage sites as a complex (**Fig. 6**). Intriguingly, work by Yang et al. (*50*) reported the recruitment of SPINDOC to DNA damage and its interaction with DDR protein, PARP1. PARP1 was also captured by SCAP for WDR76/SPIN1 (**Fig. 7A**). The recruitment of WDR76, SPIN1 and SPINDOC to DNA damages and their interactions with classic DDR proteins implied a potential model in which SPIN1 switches between complexes at the damaged site in coordination with other DDR proteins. Although, the mechanisms need to be further investigated in future studies and whether the binding or releasing of histone marks by SPIN1 complexes is related to DDR remains unknown.

The modified version of SCAP described in this work provides an upgraded option for our SCAP pipeline. Even though it might take more effort to delete a gene, the modified SCAP system could still be implemented in medium throughput studies when taking advantage of the well-established CRISPR-Cas9 system to rapidly edit the genome (*51*). Especially when using SCAP to systematically investigate different subpopulations of a protein of interest (POI), the POI will be set as the second bait, and then stable cell lines expressing the POI and a variety of 1^st^ bait proteins can be all derived from one parental cell line, in which the endogenous POI is depleted. The original SCAP workflow was easy to streamline, while the modified SCAP procedure enabled more effective enrichment. We recommend using the first version of SCAP method for rapid protein identification and using modified SCAP for more comprehensive protein complex analysis.

## Materials and Methods

### Critical Reagents, Plasmids and Cell Lines

The same critical reagents that are commercially available has been listed in (*1*). Sequences of WDR76, SPIN1 and XRCC5 open reading frame (ORF) was obtained from Kazusa Genome Technology (Kisarazu, Chiba, Japan). Vectors used for expressing Halo- or SNAP-tagged proteins, including the dual expression vector, were also described before in (*1*).

The WDR76 knockout cell line was derived from Flp-In™-293 Cell Line purchased from Invitrogen (Carlsbad, CA, USA). The knockout was achieved using RNA-guided CRISPR (Clustered regularly interspaced short palindromic repeats)-Cas9 system (*51*). Two guide RNAs were used at the same time to target the following sequences in genome.

Guide 1: 5’-GGTCGGGCGCGGCGGCTGAG-3’

Guide 2: 5’-AAGAGGGTGCATTCGTTTGG-3’

About 1 kb upstream of guide 1cutting site and about 3 kb downstream of guide 2 cutting site were cloned as homolog arms. The coding sequence of GFP was inserted between two homolog arms as a positive selection marker. In this design, the complete ORF of WDR76 is removed from genome but the endogenous promoter region is preserved to drive the expression of GFP.

An RFP expression cassette is placed in reverse direction of GFP as a positive marker for transfection and a negative marker to rule out clones that have random integration of the donor plasmid. Single cells were sorted by flow cytometry and expanded individually for screening. To screen and validate WDR76-/- clones, two pairs of primers targeting exon2 and exon13 of WDR76 were used for PCR from genomic DNA extracted from each clone.

Cell lines that stably expressing Halo-WDR76 or Halo-SPIN1 have been used in previous publications (*1, 9*). The generation of stable expression cell lines in WDR76 knockout background was similar to the stable transfection of Flp-In™-293 Cell Line described by manual (Invitrogen). All cell lines were maintained in DMEM medium with GlutaMAX, 10% FBS and 100 ug/mL hygromycin B.

Anti-SPIN1 antibodies (ab118784) are purchased from Abcam (Boston, MA, USA) and applied at 1:2000 dilution for Western blot. Anti-H3K4me3 antibodies were purchased from EMD Millipore and applied at 1:5000 dilution for Western blot.

### Single-bait Affinity Purification (AP) and Serial Capture Affinity Purification (SCAP)

All purifications were performed using stable expression cell lines, except that blank control purifications were performed on cells without expression of tagged proteins. Halo purifications were performed according to the manual of HaloTag® Mammalian Pull-Down Systems (Promega, Madison, WI) except for cells were lysed with high salt buffer (20mM Hepes at pH 7.9, 560mM NaCl, 10mM KCl, 1.5mM MgCl2, 1% Triton® X-100, 1mM Dithiothreitol, Salt Active Nucleases and Protease Inhibitor Cocktail) at a 1:3 ratio by volume (final 420mM NaCl).

For each replicate, cells were collected from 2 confluent 150mm plates. For each type of purification, three replicates were performed.

The optimized SCAP protocol is similar to (*1*). For each replicate, cells were collected from 6 confluent 150mm plates and lysed with High Salt Lysis Buffer. The lysate was centrifuged, and supernatant was collected, adjusted to 300mM of NaCl, and then incubated with SNAP-Capture Magnetic Beads (New England BioLabs, Ipswich, MA) at 4°C for 3hr. Beads were washed. Bound proteins were eluted with Elution buffer (containing PreScission Protease) at 4°C for 1hr. 10% of the eluate was aliquoted as E1 and the rest 90% was subjected to further Halo purification step. Eluate obtained above was then incubated with Magne® HaloTag® Beads (Promega) at 4°C overnight. Unbound supernatant was collected as UB2. Beads were washed. Bound proteins were eluted with Elution buffer (containing TEV protease) at room temperature for 1hr. The eluate was collected as E2. Three replicates were performed.

For comparing the pulldown of H3K4me3 by WDR76/SPIN1 and SPIN1/SPINDOC, SCAP was performed as previously described with minor adjustments using Flp-In™-293 cells stably expressing Halo-WDR76/SNAP-SPIN1 or Halo-SPIN1/SNAP-SPINDOC. SCAP enrichment was performed as previously described using 293FRT cells stably expressing a pair of baits expressing either the SNAP or the Halo tag. First enrichment was performed using SNAP beads. The eluted fractions from first enrichment were subjected to a second enrichment using Halo-ligand conjugated beads. Briefly, cells were lysed by incubating them in a high salt lysis buffer (20 mM HEPES pH 7.5, 560 mM NaCl, 10 mM NaCl, 10 mM KCl, 1.5 mM MgCl_2_, 1% v/v Triton-X 100, salt active nuclease, 1 mM DTT, Protease Inhibitor Cocktail). The lysate was centrifuged at 20,000g for 30 mins at 4°C. The supernatant was collected and incubated with SNAP beads (New England BioLabs) at 4°C. After a series of washes (20 mM HEPES pH 7.5, 0.05% v/v IGEPAL®, 1 mM DTT 300 mM NaCl, 10 mM KCl) the bait and the associated complexes were eluted from the SNAP beads by incubation with a buffer containing PreScission Protease® (20mM HEPES pH7.5, 300mM NaCl, 1mM DTT, 1mM EDTA). The eluted fraction thus obtained was next incubated with Magne HaloTag Beads (Promega) at 4°C. After repeating a series of washes, elution was performed with incubation at 4°C in a buffer containing the AcTEV protease.

The purifications of both complexes were performed in parallel and the input and E2 samples were resolved on the same gel by SDS-PAGE. The gel was then transferred and analyzed with western blot using anti-Spindlin1 (Abcam, 1:2000 dilution) and anti-H3K4me3 (EMD Millipore, 1:5000 dilution). The abundance of the SPIN1 was presented to show that comparable amount of material was used for each purification and loaded on the gel for measuring H3K4me3 pulldown. This experiment was repeated two more times yielding same results.

### Crosslinking Analysis of Halo-WDR76 Affinity Purification (AP-XL) and Halo-WDR76/SNAP-SPIN1 Serial Capture Affinity Purification (SCAP-XL)

The cells used for Halo-WDR76 AP-XL were deleted of endogenous WDR76. For each purification, a cell pellet of approximate 0.5ml was subjected to Halo purification. The Halo purification method was described in the specific method section above. Samples from 4 replicates were analyzed with mass spectrometry. The SCAP-XL of Halo-WDR76 and SNAP-SPIN1 were performed in two different scales, either with a cell pellet of ∼1.5ml or ∼3ml. The optimized SCAP protocol described in the previous method section was used in the SCAP-XL of Halo-WDR76/SNAP-SPIN1. For each scale, 3 replicates were performed. Total 6 samples were analyzed with mass spectrometry. For crosslinking reactions, eluted co-purified proteins from different purifications were crosslinked with 5mM DSSO (disuccinimidyl sulfoxide) purchased from Thermo Fisher Scientific (Waltham, MA, USA) at room temperature for 1hr. The crosslinking reaction was quenched by 50mM Tris-HCl.

### Mass Spectrometry Data Acquisition

For each sample, 5% of the amount processed for mass spectrometry analysis was loaded for SDS-PAGE and silver staining analysis first. Both uncrosslinked and crosslinked samples were processed in same procedures described before (*52*). Each sample was first TCA precipitated and then resuspended with 8M Urea buffer (in 100mM Tris-HCl, pH 8.5). The resuspended proteins were reduced with 5mM tris(2-carboxyethyl)phosphine (TCEP) and treated with 10mM 2-Chloroacetamide (CAM). Then proteins were digested with Lys-C for at least 6 hours followed dilution to 2M urea and overnight trypsin digestion. Last, the digested samples were quenched with 5% formic acid.

Multidimensional Protein Identification Technology (MudPIT) has been described before (*53*). Each digested sample was loaded onto a three-phase column. The column was pulled from capillary (100 µm i.d.) to a 5 µm tip, then packed first with 8 cm of 5 µm C18 RP particles (Aqua, Phenomenex), followed by 3.5 cm of 5 µm Luna SCX (Phenomenex), and last with 2.5 cm of 5 µm Aqua C18. Then the loaded column was washed with Buffer A (5% Acetonitrile, 0.1% Formic Acid) before placed onto instruments. All samples were analyzed using a 10-step MudPIT sequence. Each loaded column was placed in line with an Agilent 1200 quaternary HPLC pump (Palo Alto, CA) and a Velos Orbitrap Elite mass spectrometer (Thermo Scientific, San Jose, CA). MS spray voltage set at 2.5 kV; MS transfer tube temperature set at 275°C; 50 ms MS1 injection time; 1 MS1 microscan; MS1 data acquired in profile mode; 15 MS2 dependent scans; 1 MS2 microscan; and MS2 data acquired in centroid mode. MS1 scans acquired in Orbitrap (OT) at 60000 resolution; full MS1 range acquired from 400 to 1500 m/z; MS1 AGC targets set to 1.00E+06; MS1 charge states between 2-5; MS1 repeat counts of 2; MS1 dynamic exclusion durations of 90 sec; ddMS2 acquired in IT; MS2 collision energy and fragmentation: 35% CID; MS2 AGC targets of 1.00E+05; MS2 max injection times of 150 ms.

The LC-MS method for analyzing crosslinked samples has been described before (*1*). Crosslinked peptides were analyzed on an Orbitrap Fusion Lumos mass spectrometer (Thermo Scientific, San Jose, CA) coupled to a Dionex UltiMate 3000 RSCLnano System. Peptides were loaded on the Acclaim™ PepMap™ 100 C18 0.3 mm i.D. 5 mm length trap cartridge (Thermo Scientific, San Jose, CA) with loading pump at 2 µl/min via autosampler. Analytical column with 50 µm i.D. 150 mm length, was packed in-house with ReproSil-Pur C18-AQ 1.9 μm resin (Dr. Masch GmbH, Germany). The organic solvent solutions were water/acetonitrile/formic acid at 95:5:0.1 (v/v/v) for buffer A (pH 2.6), and at 20:80:0.1 (v/v/v) for buffer B. When peptides were analyzed, the chromatography gradient was a 20 min column equilibration step in 2% B; a 10 min ramp to reach 10% B; 120 min from 10 to 40 % B; 5 min to reach 95% B; a 14 min wash at 95% B; 1 min to 2% B; followed by a 10 min column re-equilibration step at 2% B. The nano pump flow rate was at 180 nL/min. An MS3 method was made specifically for the analysis of DSSO cross-linked peptides. Full MS scans were performed at 60,000 m/z resolution in the orbitrap with 1.6 m/z isolation window, and the scan range was 375-1500 m/z. Top 3 peptides with charge state 4 to 8 were selected for MS2 fragmentation with 20% CID energy. MS2 scans were detected in orbitrap with 30,000 m/z resolution and dynamic exclusion time is 40 s. Among MS2 fragments, if two peptides with exactly mass difference of 31.9720 with 20 ppm mass tolerance, both were selected for MS3 fragmentation at CID energy 35% respectively. MS3 scans were performed in the ion trap at rapid scan with isolation window of 3 m/z, maximum ion injection time was 200 ms. Each MS2 scan was followed by maximum 4 MS3 scans. For the analysis of crosslinked samples in this work, a FAIMS pro module (Thermo Scientific, San Jose, CA) was installed, and compensation voltage (CV) was set to alternate between -40, -60 and - 80V during acquisition.

### MudPIT Data Analysis

Collected MS/MS spectra were searched with the ProLuCID (*54*) software against a database of 163860 protein sequences combining 81592 non-redundant Homo sapiens proteins (NCBI, 2020-11-23 release), 426 common contaminants, and their corresponding 81930 randomized amino acid sequences. In this manuscript, sequences of Halo tag, SNAP tag, AcTEV protease and PreScission Protease were added to contaminants. All cysteines were considered as fully carboxamidomethylated (+57 Da statically added), while methionine oxidation was searched as a differential modification. The following dynamic modifications were included during analysis: mono-/di-/tri-methylation and acetylation of lysine; mono-/di-methylation of arginine. DTASelect (*55*) v1.9 and swallow, an in-house developed software, were used to filter ProLuCID search results at given FDRs at the spectrum, peptide, and protein levels. Here, all controlled FDRs were less than 1%. Data sets generated from each experiment were contrasted against their merged data set using Contrast v1.9 and in house developed sandmartin v0.0.1. Our in-house developed software, NSAF7 v0.0.1, was used to generate spectral count-based label free quantitation (*56*). The contaminants and keratin proteins were eliminated from the total. The dNSAF values generated by NSAF7 were defined in (*56*). In short, dNSAF stands for normalized spectral abundance factor (NSAF) computed using distributed spectra counts. For plotting the abundance of histone H2A, H2B and H3, distributed spectra among variants of each type of histone were summed. To compare datasets, QSPEC (*41*) was used to determine whether the abundances of a protein in two samples were statistically significant. Log fold change, FDRs and Z score (Zstatistic) was generated by QSPEC. Only proteins have QSPEC FDR<0.05 and Z score value>2 were considered significantly enriched in one sample over another (or Z score<-2 for proteins enriched in the later sample than in the former one) (**Data S2F**). Full dNSAF tables with QSPEC results can be found in **Data S2A-C**. To compare the Halo-WDR76/SNAP-SPIN1 SCAP-MS data to published Halo-SPIN1/SNAP-SPINDOC SCAP-MS data (*1*) (**Data S2D**), the new datasets were also analyzed using the same pipeline. The analysis method was the same as described above except for the use of a different database (Homo sapiens proteins, NCBI, 2016-06-10 release). For the post translational modifications of histone H3, the percentages of modified H3K9 were calculated from the spectral counts of peptides containing modified H3K9 and total H3K9-containing peptides (**Data S2E**). To confirm the result, we also calculated the percentages using the intensity of peptides, the results is consistent with the spectral counts based quantification method.

### Crosslinking Data Analysis

For the data analysis of DSSO cross-linked peptides Proteome Discoverer 2.4 (Thermo Scientific, San Jose, CA) with add on XLinkX node was used in peptide identification and cross-linked peptide searching. The following settings were used: precursor ion mass tolerance, 10 ppm; fragment ion mass tolerance, 0.6 Da; fixed modification, Cys carbamidomethylation; variable modification, Met oxidation, Lys DSSO Amidated, and Lys DSSO hydrolyzed; maximum equal dynamic modification, 3. Proteins FDR was set at 0.01. Acetylation was selected as a dynamic modification. The search engine files for crosslinking datasets are available at MassIVE (**Data S1A**). The crosslinking results were visualized in 2D by a web based software xiView (*42*) (https://xiview.org/xiNET_website/index.php). Tables uploaded to xiView can be found in **Data S3**.

### Imaging Sample Preparation

All cells used for imaging were plated into Mat-Tek dishes with No 1.5 coverslip bottoms. Imaging samples were kept in phenol red free DMEM Medium with GlutaMAX and 10% FBS. While imaging, live cells are kept under 37°C, 5% CO2 and humidified condition. For WDR76 and SPIN1 imaging experiments, pcDNA5FRT-SNAP-SPIN1 or pcDNA5FRT-SNAP were transfected to Halo-WDR76 stable expression cell line (with endogenous WDR76). For negative control experiments, pcDNA5FRT-SNAP was co-transfected with pcDNA5FRT-Halo to the original Flp-In™-293 Cell Line (Invitrogen); and for positive control experiments, pcDNA5FRT-Halo-NLS-SNAP was transfected. For microirradiation assay, SNAP tagged proteins were transiently expressed in Halo-WDR76 stable expression cell line (with endogenous WDR76). Cells were imaged 24hr after transfection.

Halo tags were stained with HaloTag® Ligands TMRDirect (Promega), and SNAP tags were stained with SNAP-Cell® 505-Star (NEB). HaloTag® Ligands TMRDirect were added to medium to a final concentration of 50nM and incubated overnight. SNAP-Cell® 505-Star ligands were added to medium at the same day of imaging, then cells were incubated with the ligands and washed. For microirradiation assay, HOECHST 33258 was also added together with SNAP ligands at a final concentration of 5μg/ml. After staining, cells were allowed to stabilize in fresh medium for 30min in the incubator before imaging.

### Acceptor Photobleaching Förster Resonance Energy Transfer (apFRET)

apFRET was performed similarly to Weems et al. (*57*). In detail, data was acquired with a PerkinElmer Life Sciences UltraVIEW VoX spinning disk microscope controlled by Velocity software. The microscope is equipped with Yokogawa CSU-X1 Spinning disk scanner, ORCA-R2 camera (Hamamatsu C10600-10B), EMCCD (Hamamatsu C9100-23B) and PhotoKinesis accessory. The base of the microscope is Carl Zeiss Axiovert 200M. The Dichroic passes 405, 488, 561 and 640nm. HaloTag® Ligands TMRDirect was excited by 561nm laser and collected through 445 (W60), 615 (W70) filter. SNAP-Cell® 505-Star was excited by 488nm laser and collected through 525 (W50) filter. To collect AP-FRET data, time lapse movies were recorded to collect at least 10 timepoints before and after accepter photobleaching. The movies were recorded at a speed of one image per second. Images were recorded using ORCA-R2, and objective was 40x (Oil, NA=1.3).

For each cell, accepters were bleached by 561nm laser at 100% laser power for 10 cycles. The donor intensity before (I_Before_) and after (I_After_) were, separately, averaged over time. FRET efficiency (E) was represented as: E=1- (I_Before_ / I_After_). E values were calculated in batch using in house imageJ (National Institutes of Health) plugin (accpb FRET analysis jru v1). For statistically comparing the FRET efficiency between different protein pairs, the measurements for each pair were first tested for normal distribution, and then were compared using two-tailed t-test in OriginPro (OriginLab, Northampton, MA). Measurements and statistical results can be found in **Data S5A**.

### Fluorescence Cross-Correlation Spectroscopy (FCCS)

FCCS data was acquired using a ConfoCor 3 with an LSM-510 (Zeiss) microscope. Cells were imaged through a C-Apochromat 40x (NA=1.2) objective. The coverslip thickness correction collar was adjusted to achieve maximum signal in the first Mat-Tek dish. In all cases, the pinholes for all channels were aligned just before acquisition. It was left at that position for the rest of the acquisitions. SNAP-Cell® 505-Star was excited at 488nm, and its fluorescence was collected through a 505-540 bandpass filter. HaloTag® Ligands TMRDirect was excited at 561nm, and its fluorescence was collected through a 580 longpass filter. To prevent cross-talk between the fluorophores, Pulsed Interleaved Excitation (PIE) laser switching was employed. The 488 and 561nm lasers were alternately switched on for 100μs total period (50μs for each laser) with a 2.5μs illumination delay. For each pair of proteins, 10 curves were measured and used for data analysis. Files were analyzed in Fiji (https://fiji.sc/) using in-house written plugins (analysis cross corr jru v2). More detailed results from the analysis can be found in **Data S5B**.

### UV-Laser Microirradiation

UV-laser microirradiation assay was also performed on UltraVIEW VoX spinning disk microscope. The excitation and emission filter settings for each fluorophore has been stated above. Time lapse movies were recorded using ORCA-R2 at speed of one image per second. Objective was 40x (Oil, NA=1.3). The assay has been used in Gilmore et al. (*9*) to test the recruitment of Halo-WDR76. Similarly, each cell was irradiated in a 1 pixel (∼0.17μm) width stripe area near the center of the nucleus. The microirradiation was induced with 405nm laser at 100% laser power (20 cycles). The fluorescence intensities at each timepoint (t) of the stripe (It) and the whole nucleus (Tt) were measured cell by cell using imageJ. The timepoint starting microirradiation was defined as t=0. Recruitment (Rt) was defined as: Rt=(It/Tt)/(I0/T0). For each experiment, multiple cells were measured and averaged. Rt values were calculated in batch using in-house written plugin (dna damage analysis multicolor jru v1). The highest RT (MaxRT) values were also obtained from the plugin. Detailed values of RT and MaxRT are available in **Data S5C and D**.

To compare the kinetics of the recruitments, the averaged RT values from timepoint 0 to 60 of Halo-WDR76/SNAP-SPIN1 and Halo-WDR76/SNAP-XRCC5 were fitted to a modified Hill function in OriginPro (OriginLab, Northampton, MA):

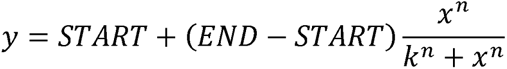

Here, START stands for the y value at the beginning of the curve and END stands for the y value when it reaches the maximum and the curve turns flat. k is the Michaelis constant and n is the Hill coefficient. The time for the recruitment of each protein taken to reach its half MaxRT (t_MaxRT/2_) was used to estimate the rate of recruitment. t_MaxRT/2_ values were obtained either directly from the k values provided by OriginPro or manually measured from the fitted curves. Fitting results can be found in **Data S5E**.

### Structure Prediction and Integrative Complex Modeling

All 3D structures were visualized and manipulated by UCSF Chimera (*58*) to generate figures. Distances between the α-carbons of crosslinked lysine residues (Cα-Cα) were also measured in Chimera.

For structure predictions, the full length models of WDR76 (Uniprot: Q9H967) and SPIN1 (Uniprot: Q9Y657) were generated by I-Tasser (*43, 44, 59*) On-line Server (https://zhanglab.ccmb.med.umich.edu/I-TASSER/) based on protein sequences. We next measured the distances between each pair of crosslinked lysine residues in every model (**Data S4A**). We allowed the maximum Cα-Cα distance of 35 Å between DSSO-crosslinked lysine residues to be considered satisfied. For WDR76 models, we found both model3 and model5 were reasonably folded and satisfied 5 out of 6 intra-WDR76 crosslinks (**Fig. S3A**). For SPIN1, models are similar (**Fig. S3B**) and we picked model2 as our best model for later analysis as it satisfied all 7 intra-SPIN1 crosslinks.

We next performed interaction space analysis using the quick scanning mode of DisVis (*45, 46*) (https://wenmr.science.uu.nl/disvis/). Either WDR76 model3 or model5 were used as the fixed chain and SPIN1 model2 was used as the scanning chain. The 3 crosslinks detected between WDR76 and SPIN1 by SCAP-XL were used as restraints. For the analysis using WDR76 model3, no restraint was reported as false positive by DISVIS. For WDR76 model5, one restraint out of the 3 had a Z-score over 1 and suggested to be false positive by DISVIS (**Data S4B**). We therefore picked WDR76 model3 for the WDR76/SPIN1 complex modeling analysis.

Molecular docking was performed using HADDOCK 2.4 webserver (*47, 48*) (https://wenmr.science.uu.nl/haddock2.4/). The docking was performed following HADDOCK tutorials and the protocol described by de Vries et al. (*47*) Inter crosslinks of WDR76 and SPIN1 were used as unambiguous restraints. The center of mass restraints option was switched on for all docking calculations. We used the same sampling and clustering parameters for all analyses. HADDOCK performed three steps for docking. (*60*) First, randomization and rigid body energy minimization were performed (“it0”). 180 degrees rotated solutions were sampled during the rigid body energy minimization step. Up to 10,000 models were sampled and 1,000 models were saved at this stage. The best 200 models were subjected to semi-flexible refinements (“it1”), then refined in water as the final step (“water”). The refined models were clustered using the Fraction of Common Contacts (FCC) (*61*) method with a fraction of contacts cutoff of 0.6. For the docking of WDR76 model3 and SPIN1 model2, HADDOCK clustered 127 structures in 13 clusters and reported the best 4 structures for the top 10 clusters in their result page (**Data S4C**). RMSD values were provided in the downloaded result package from HADDOCK. The average RMSD values were calculated by cluster for either all the structures or the best 4 structures in each cluster. The rmsd values were computed by comparing to the best structure in each cluster; the rmsd-Emin values were computed by comparing to the structure with the overall lowest energy. From the HADDOCK stats, cluster6 has the lowest HADDOCK score and this score is significantly better than the next cluster. The best 4 structures in cluster6 have a small rmsd of 1.3Å. The Cα-Cα distances between crosslinked lysine residues in the best 4 structures were measured (**Data S4D**). Using 35 Å as a cutoff, all 3 crosslinks are satisfied by these 4 structures. Therefore, we picked the structure cluster6-1 with the lowest HADDOCK score as our best model for visualization.

## Supporting information

Supplementary Materials - Figures and DataSet Descriptions

DataS1

DataS2

DataS3

DataS4

DataS5

## Acknowledgments

## Funding

National Institutes of Health grant R35GM118068 (JLW)

National Institutes of Health grant T32GM138077 (JC)

National Institutes of Health grant RO1GM112639 (MPW)

National Institutes of Health grant R35GM145240 (MPW)

Stowers Institute for Medical Research (JLW, MPW)

## Author Contributions

Conceptualization: XL, YZ, JLW, LF, MPW

Investigation: XL, YZ, ZW, YH, CASB, JL, JC, SB, BDS, JRU

Formal Analysis: XL, YZ, ZW, YH, CASB, JL, JC, SB, BDS, JRU, LF

Resources: XL, YZ, YH, JC, SB

Software: ZW, BDS, JRU, LF

Data Curation: ZW, BDS, JRU, LF

Writing: XL, MPW

Supervision: LF, JLW, MPW

Funding acquisition: JLW, MPW

## Competing Interests

The authors declare no competing financial or non-financial interests.

## Data and Materials availability

The availability of mass spectrometry datasets in public repositories is provided in Data S1 and Data S3. Additional Data for the analysis and processing of all mass spectrometry data is provided in Data S2 and Data S4. The data regarding the fluorescent imaging experiments are provided in Data S5. Materials inquiry requests should be made to.

## Notes

### Competing Interest Statement

The authors have declared no competing interest.

