## Supplementary Materials - Figures and DataSet Descriptions for "Serial Capture Affinity Purification and Integrated Structural Modeling of the H3K4me3 Binding and DNA Damage Related WDR76:SPIN1 Complex"

**This PDF file includes:**

Figs. S1 to S5

Data S1 to S5

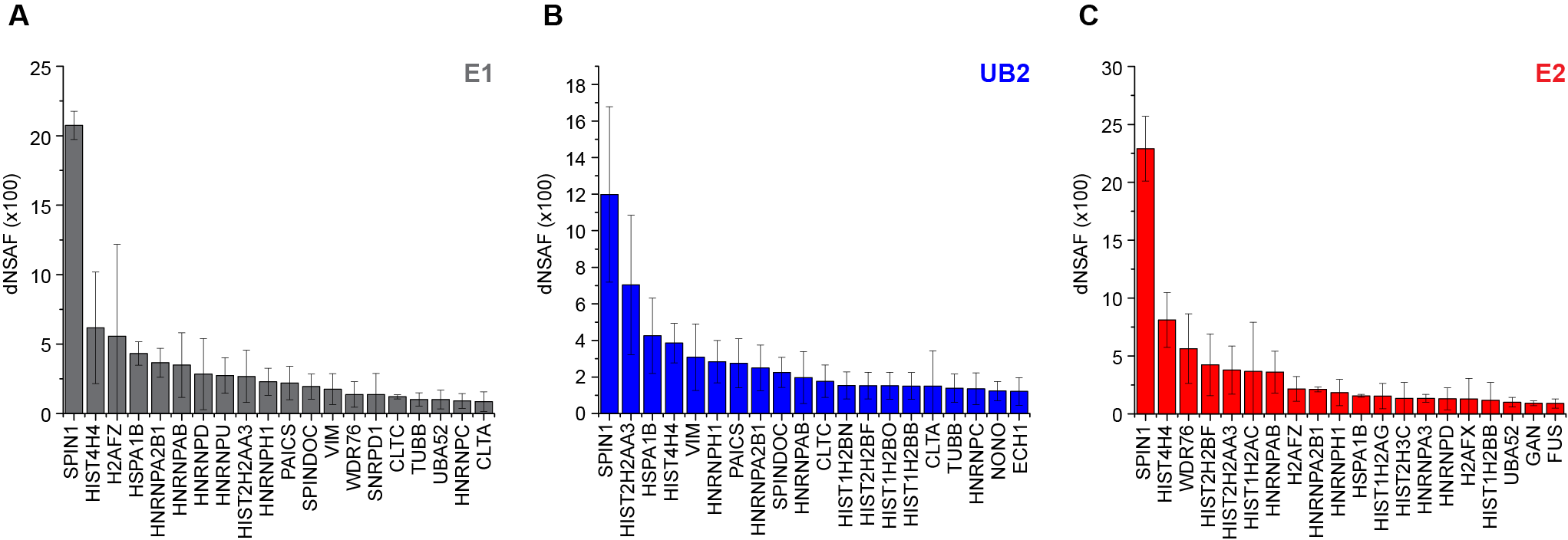

**Fig. S1. Top 20 proteins ranked by abundance identified by MudPIT in E1 (A), UB2 (B), and E2 (C) fractions of SCAP using Halo-WDR76 and SNAP-SPIN1 as bait proteins.** Each dNSAF value plotted is the average from three biological replicates; error bars represent standard deviations.

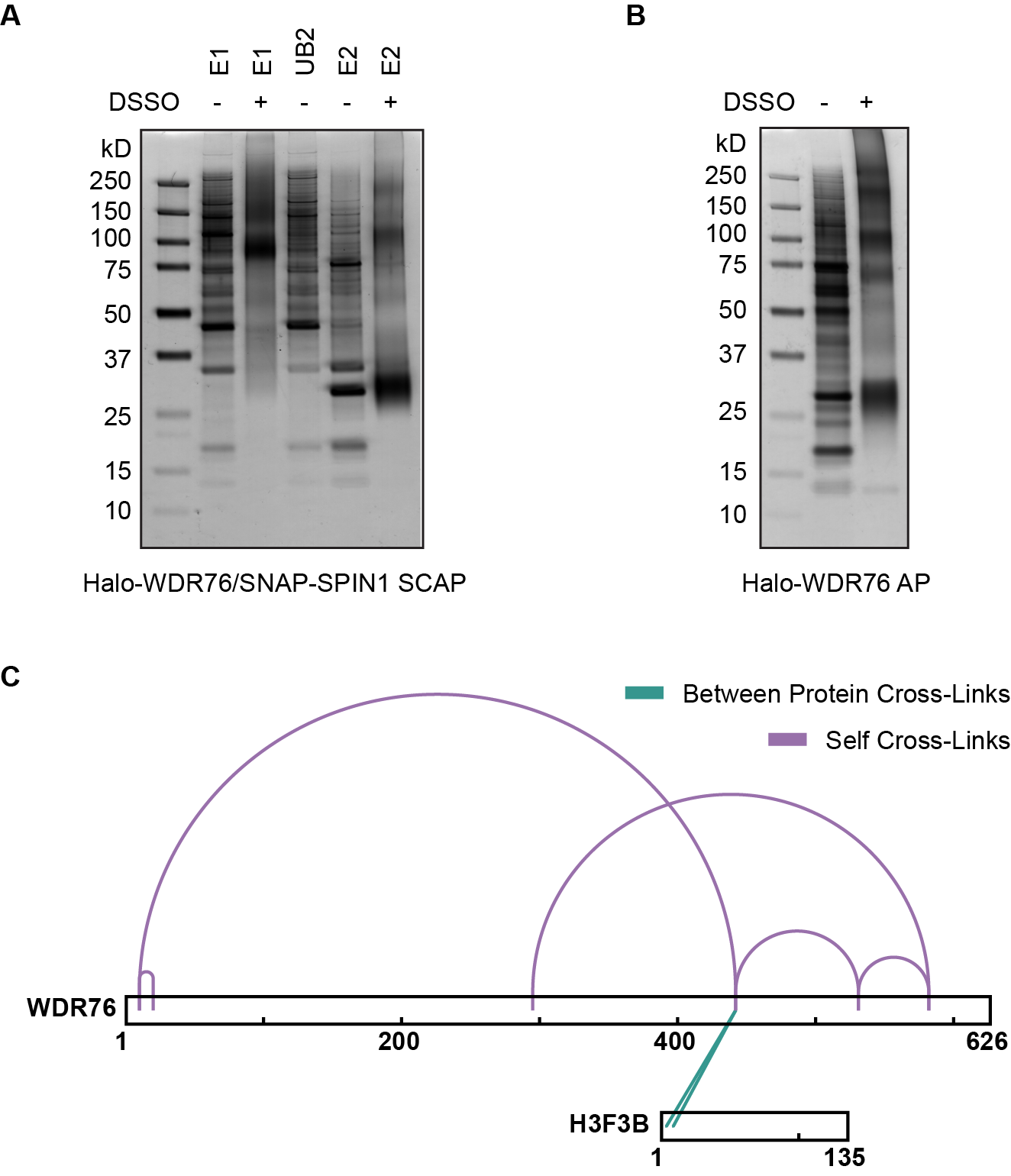

**Fig. S2. Gel analysis of DSSO crosslinked samples and crosslinking mass spectrometry analysis of Halo-WDR76 single-bait purification.** (**A**) Halo-WDR76/SNAP-SPIN1 SCAP purification and crosslinking. SNAP-SPIN1 was first purified followed by Halo-WDR76. E1 and E2 samples were crosslinked with DSSO. Samples were analyzed by SDS-PAGE gel and silver staining. 5% of crosslinked E2 sample used for mass spectrometry analysis were loaded. (**B**) Halo-WDR76 single-bait affinity purification and crosslinking. Co-purified proteins with or without DSSO crosslinking were analyzed by SDS-PAGE gel and silver staining. 5% of crosslinked sample used for mass spectrometry analysis were loaded. Same amount of protein standards was used in panel **A** and **B**. (**C**) DSSO crosslinked Halo-WDR76 purification samples analyzed by mass spectrometry. Linear view of protein-protein interactions that are linked by at least 2 crosslinks by xiView (*42*).

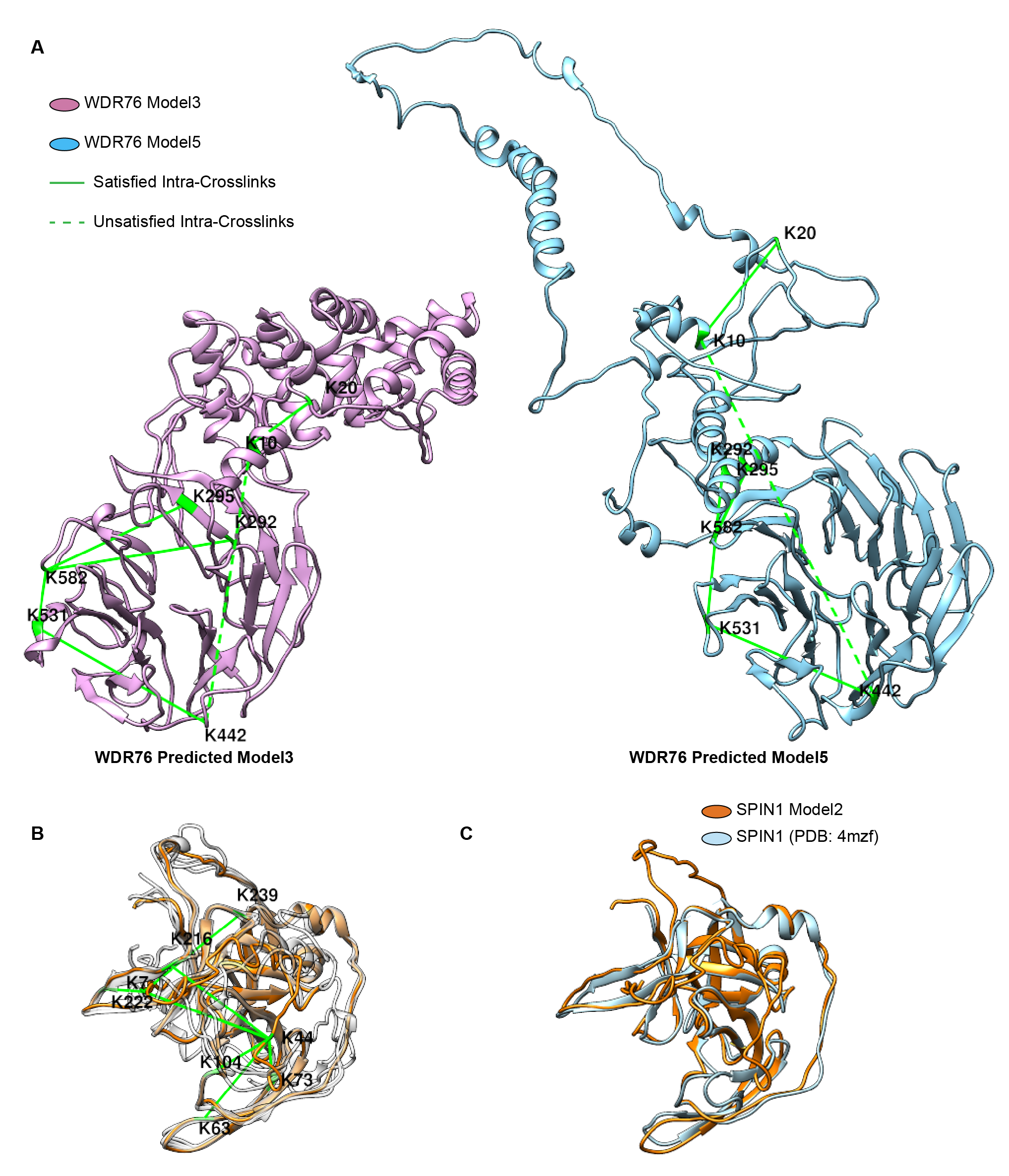

**Fig. S3. Prediction of full length WDR76 and SPIN1 structures.** (**A**) Ab initio prediction of full length WDR76 structure. Structural models were predicted by I-TASSER (*44*) from amino acid sequence. Identified self-crosslinks of WDR76 were mapped to each model and Cα-Cα distances between crosslinked lysine residuals were measured. The best two models, model3 and model5, were displayed with crosslinked lysine labeled and colored in green. Crosslinks are visualized as green lines, while the unsatisfied crosslinks (with Cα-Cα distance between crosslinked sites larger than 35Å) are shown in dash. (**B**) Ab initio prediction of full length SPIN1 structure. Structural models were predicted by I-TASSER (18) from amino acid sequence. Total five models are aligned and displayed in overlays. Model2, which is the best model to satisfy all the identified self-crosslinks of SPIN1, is colored in orange. Crosslinked sites were labeled and colored in green. Crosslinks are visualized as green lines. (**C**) The best predicted full length SPIN1 structure model (colored in orange) is aligned and compared to published SPIN1 structure (chain B of PDB: 4mzf, colored in light blue).

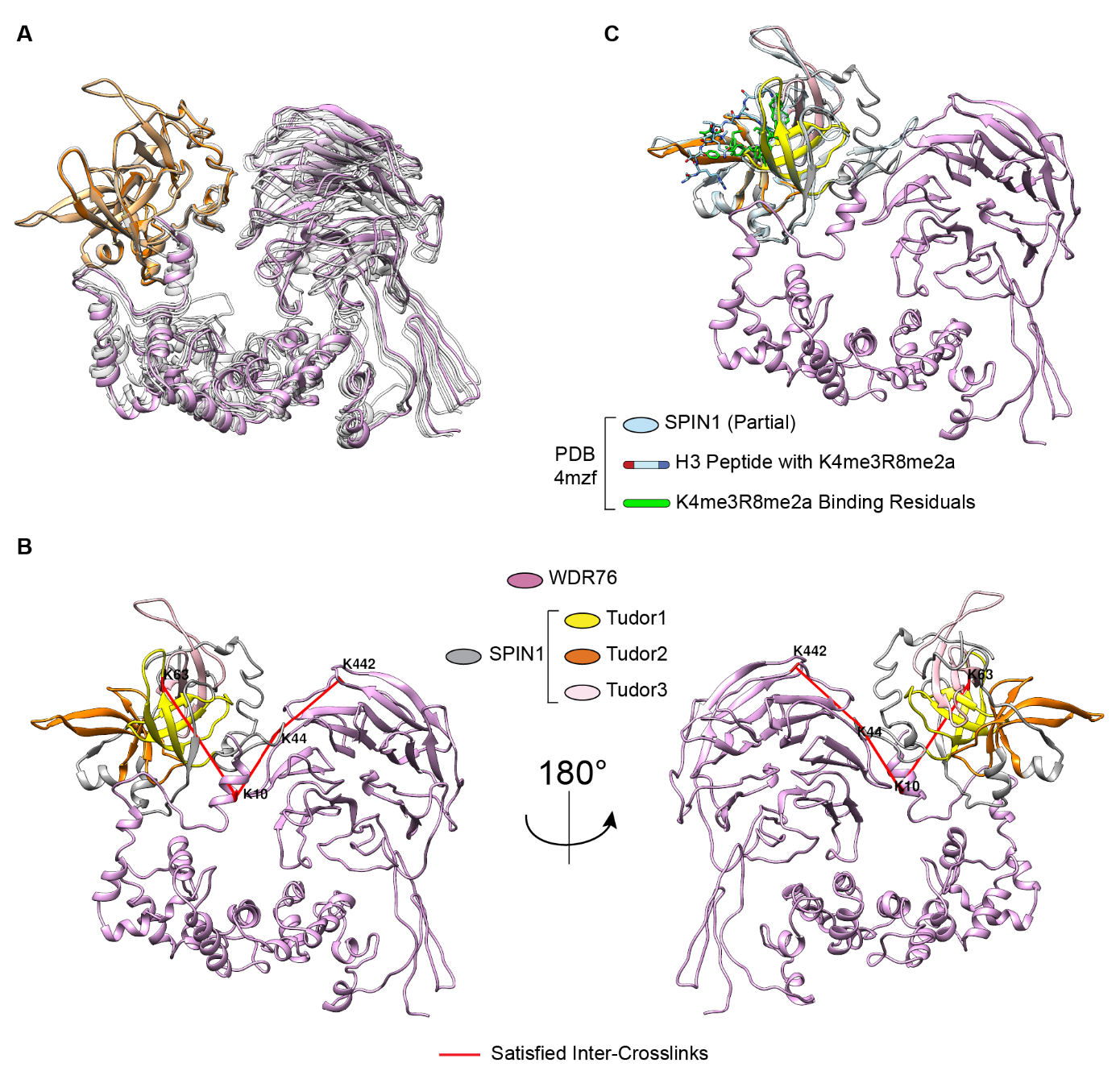

**Fig. S4. Structure modeling of WDR76/SPIN1 complex.** (**A**) Docking of predicted model3 of WDR76 and model2 of SPIN1 with HADDOCK (*48*). Inter-crosslinks between WDR76 and SPIN1 were used in docking as restraints. An ensemble representation of the best four structures in the top cluster with the lowest HADDOCK score is displayed. The structure in solid color (WDR76 in plum color and SPIN1 in orange) has the best HADDOCK score in the cluster. (**B**) Visualization of crosslinks between WDR76 and SPIN1 in the heterodimer model with the best HADDOCK score. The three Tudor domains of SPIN1 are color coded. Crosslinked sites were labeled and colored in red. Crosslinks are visualized as red lines. (**C**) Visualization of the heterodimer model of WDR76/SPIN1 complex with the best HADDOCK score. A known structure of SPIN1 Tudor domains (light blue) containing a modified histone H3 N’-peptide (PDB: 4mzf) is aligned to the model. Key residuals recognizing the modifications of H3 are colored in green.

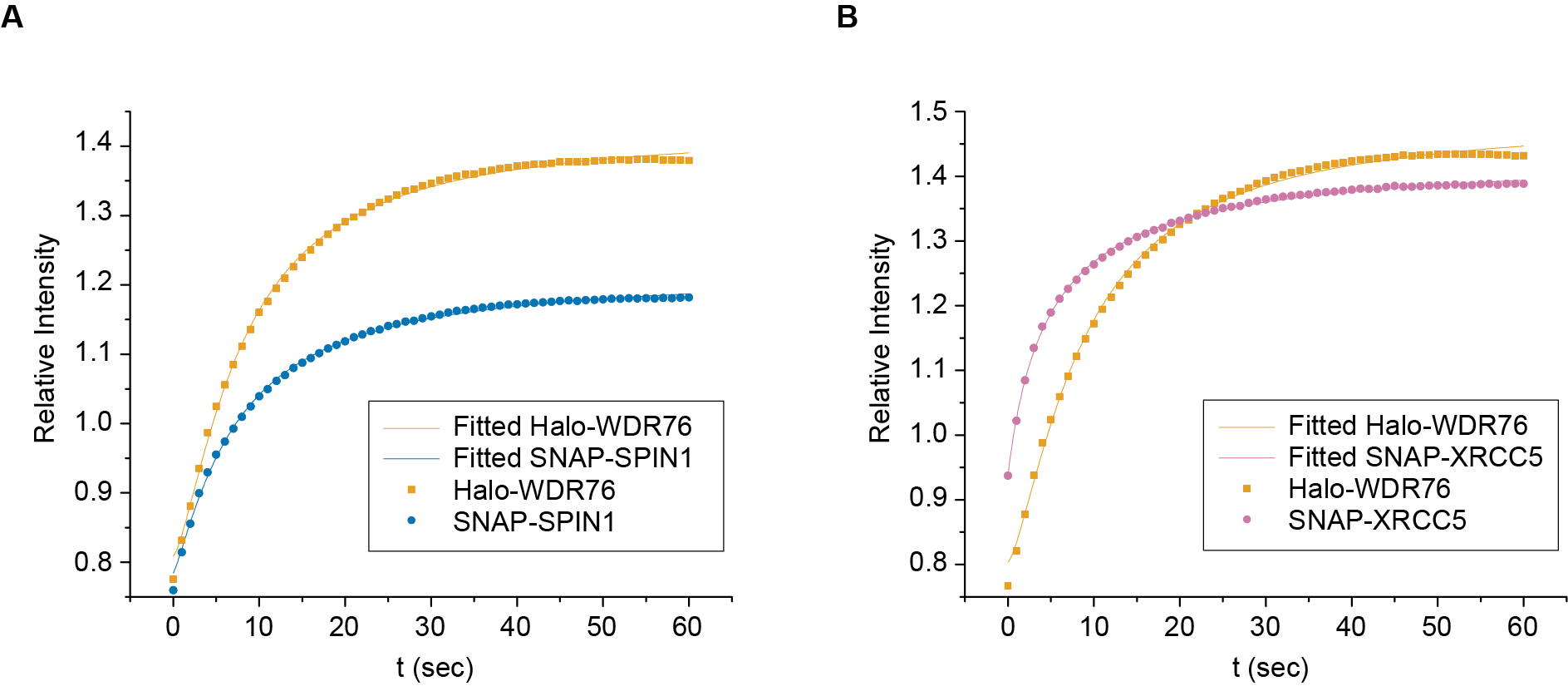

**Fig. S5. The co-recruitment of WDR76 and SPIN1 to microirradiated region.** The relative intensities of each protein at the laser-irradiated stripe were recorded for each second. The measurements from the moment of microirradiation up to 1 minute were used for nonlinear fitting in OriginPro (OriginLab, Northampton, MA). Each data point is an average from all the measured cells. (**A**) Halo-WDR76 and SNAP-SPIN1 were co-expressed and measured. (**B**) Halo-WDR76 and SNAP-XRCC5 were co-expressed and measured.

**Data S1. Information of mass spectrometry datasets.** (**A**) Full mass spectrometry data accessibility. (**B**) Spectra and peptide FDR information of MudPIT experiments. (**C**) FDR reports of crosslinking mass spectrometry datasets by Proteome Discoverer 2.4 (Thermo Scientific, San Jose, CA).

**Data S2. Additional tables of MudPIT data.** (**A**) Full contrast table of Halo-WDR76 and Halo-SPIN1 AP-MS data. (**B**) Full contrast table of Halo-WDR76/SNAP-SPIN1 SCAP-MS data. (**C**) Full contrast table of Halo-WDR76 AP-MS data generated using cells without endogenous WDR76. The data of E2 fraction in Data S2B was also compared in the same table. In all the contrast tables, peptide counts, spectral counts, sequence coverages, dNSAF values and QSPEC (*41*) results are included. (**D**) Comparing different SPIN1 containing complexes from SCAP-MS data. Data searched using 2016 database to be compared to published SPIN1/SPINDOC data (*1*). (**E**) Spectral counts of total and modified peptides of histone H3 containing K9. The percentage values were calculated from spectral count or intensity. (**F**) List of proteins enriched in SCAP E2 fraction over other samples. NCBI RefSeq protein accessions are used here to distinguish protein identities.

**Data S3. Additional tables of XL-MS uploaded to xiView** (*42*) **for visualization.** (**A**) Full table of Halo-WDR76/SNAP-SPIN1 SCAP-XL data. (**B**) Annotations of WDR76 and SPIN1. (**C**) Metadata for of Halo-WDR76/SNAP-SPIN1 SCAP-XL. (**D**) Full table of Halo-WDR76 AP-XLMS data. (**D**) Metadata for Halo-WDR76 AP-XLMS. The protein names provided in metadata are gene names used by Uniprot database.

**Data S4. Additional data of structural modeling.** (**A**) Cα-Cα distances between crosslinked sites measured from predicted WDR76 and SPIN1 models. (**B**) Results of DisVis (*45, 46*) analysis. (**C**) Parameters, statistic results and cluster RMSD values of HADDOCK analysis. (**C2**) HADDOCK statistics and RMSD values for docking of WDR76 model5 and SPIN1. (**D**) Cα-Cα distances between inter-protein crosslinked sites measured from the top WDR76/SPIN1 heterodimer models.

**Data S5. Additional data of fluorescent imaging experiments.** (**A**) Measurements and statistics of apFRET experiments. (**B**) Measurements and fitting of FCCS experiments. (**C**) Measurements of UV laser-induced microirradiation assay. (**D**) Maximum recruitments and statistics of microirradiation assay. (**D**) Non-linear fitting results of recruitments in the microirradiation assay.
